# Mapping genetic variants for nonsense-mediated mRNA decay regulation across human tissues

**DOI:** 10.1101/2022.10.19.512888

**Authors:** Bo Sun, Liang Chen

## Abstract

**Background:** Nonsense-mediated mRNA decay (NMD) was originally conceived as an mRNA surveillance mechanism to prevent the production of potentially deleterious truncated proteins. Recent research shows NMD is an important post-transcriptional gene regulation mechanism selectively targeting many non-aberrant mRNAs. However, how natural genetic variants affect NMD and modulate gene expressions remains elusive.

**Results:** Here we elucidate NMD regulation of individual genes across human tissues through genetical genomics. Genetic variants corresponding to NMD regulation are identified based on the GTEx data through unique and robust transcript expression modelling. We identify genetic variants that influence the percentage of NMD-targeted transcripts (pNMD-QTLs), as well as genetic variants regulating the decay efficiency of NMD-targeted transcripts (dNMD-QTLs). Many such variants are missed in traditional expression quantitative trait locus (eQTL) mapping. NMD-QTLs show strong tissue specificity especially in the brain. They are more likely to colocalize with disease single-nucleotide polymorphisms (SNPs). Compared to eQTLs, NMD-QTLs are more likely to be located within gene bodies and exons, especially the penultimate exons from the 3’ end. Furthermore, NMD-QTLs are more likely to be found in the binding sites of miRNAs and RNA binding proteins (RBPs).

**Conclusions:** We reveal the genome-wide landscape of genetic variants associated with NMD regulation across human tissues. Our analysis results indicate important roles of NMD in the brain. The preferential genomic positions of NMD-QTLs suggest key attributes for NMD regulation. Furthermore, the colocalization with disease-associated SNPs and post-transcriptional regulatory elements implicate regulatory roles of NMD-QTLs in disease manifestation and their interactions with other post-transcriptional regulators.

## Background

Nonsense-mediated mRNA decay (NMD) is an important post-transcriptional regulatory mechanism defining gene expression. It was originally conceived as an mRNA surveillance mechanism that recognizes and degrades transcripts harboring a premature termination codon (PTC) to prevent the production of potentially deleterious truncated proteins [1,2]. These mRNAs with PTCs originate from various sources (e.g., nonsense mutations, aberrant alternative splicing, and DNA rearrangements) [3]. The broader impact of NMD has been revealed in genetic disease, gene editing, and cancer immunotherapy [4]. Nonsense mutations account for 20.3% of all disease-associated single base-pair mutations and many of them introduce PTCs [5]. While NMD is a protective mechanism against production of C-terminal truncated proteins, NMD can either aggravate or alleviate the effects of those PTCs that cause genetic diseases [6].

Recent research shows NMD is an important post-transcriptional gene regulation mechanism targeting many non-aberrant mRNAs [7–11]. Transcriptome-wide studies suggest NMD accounts for autoregulation of 3%-10% normal transcripts in different cell types [12–14]. Our own analysis of the GENCODE annotation showed that about 12% of natural transcripts are presumably targeted by NMD regulation, since they follow the 50-nt rule (i.e., the presence of a terminal codon >50 nt upstream of the last exon-exon junction). Accordingly, NMD factors are implicated in cellular homeostasis, stress response, cell cycle, differentiation, development, neural activity, immunity, and spermatogenesis [15–24]. NMD typically downregulates its substrate by 2-20 folds [25,26], generating significant functional and clinical implications. Although the overall degradation process is believed to be similar for both erroneous and normal transcripts, factors involved in NMD recognition and regulation of degradation efficiency are poorly understood.

Variations in NMD efficiency are observed across tissues and individuals [27,28]. Such variations can influence the outcome of PTC read-through therapeutic strategies aimed at suppressing nonsense mutations to restore full-length proteins, as the efficacy of such nonsense suppression largely depends on NMD efficiency (reviewed in [29]). Thus, a thorough examination of natural genetic variants affecting NMD efficiency is essential in understanding not only NMD regulation but also phenotypic variability and different degrees of expressivity manifested in various nonsense-mutation-triggered diseases.

Genetic variants associated with changes in gene expression (expression quantitative trait loci, eQTLs) are being catalogued for human tissues on an unprecedented scale (e.g., the Genotype-Tissue Expression (GTEx) project) [30,31]. Genetical genomics has been focused on identifying genetic loci linked to variation(s) in mRNA expression levels. Recent studies have extended the analysis to identify genetic variants associated with other molecular phenotypes (e.g., ribosome occupancy, protein abundance, allele-specific expression, and alternative splicing) [32–34]. eQTLs mapped to NMD factors (e.g., *SMG7, UPF3B, MAGOH*, and so on) provide valuable information about how genetic variants may affect the NMD pathway through their cis-acting regulation on NMD factors. Several studies examined how loss-of-function variants and protein-truncating variants affect their own gene expression, which implicated the involvement of surveillance NMD [35–37]. However, no systematic survey of *cis* genetic variants on individual NMD substrates and their impact on NMD-regulation has been done.

Here we utilize the GTEx resources and build a genome-wide landscape of tissue-specific genetic variants involved in NMD regulations. To distinguish the NMD effect on steady-state mRNA levels from that of transcriptional regulation, we built a novel and robust statistical model. We focused on natural mRNAs targeted by NMD according to the 50-nt rule instead of aberrant mRNAs caused by DNA mutations or RNA processing errors. We identified natural genetic variants controlling the percentage of NMD-targeted transcripts, and variants regulating the decay efficiency of NMD-targeted transcripts.

## Results

### Modelling of NMD-QTLs

We propose a statistical model to identify and characterize genetic variants associated with NMD and apply it to human tissue data (Fig. 1a). The mRNA transcripts of a gene were divided into two categories: NMD-targeted and non-NMD transcripts, based on whether they were annotated by the “nonsense_mediated_decay” tag in the GENCODE annotation. Genes with both NMD-targeted and non-NMD transcripts were referred as NMD-genes. We used *α* and *θ* to parameterize NMD regulation: *α* represents the percent of transcribed transcripts targeted by NMD, and *θ* represents the percent of NMD-targeted transcripts remaining after degradation. As illustrated in Fig. 1b, gene haplotypes bearing different alleles (‘A’ or ‘a’) may generate different amounts of transcribed mRNAs (*t*_A_ or *t*_a_), different percentages of transcribed mRNAs (*α*_A_ or *α*_a_) may be targeted by NMD regulation, and these NMD-targeted transcripts may be degraded at different efficiencies (1−*θ*_A_ or 1−*θ*_a_). Note that degradation efficiency is relative to the regular degradation of non-NMD transcripts. Generally, genetic variants identified from a regular eQTL analysis are those with allele-specific transcription levels (i.e., *t*_A_≠*t*_a_) without taking into consideration NMD-transcripts (*α*) and their degradation (1−*θ*). Here, we aim to capture genetic variants resulting in different *α*s or *θ*s. The former, called pNMD-QTLs, controls the different <u>p</u>ercentages of NMD-targeted transcript isoforms (*α*_A_≠*α*_a_). The latter, called dNMD-QTLs, regulates the <u>d</u>egradation efficiency of NMD-targeted isoforms (*θ*_A_ ≠*θ*_a_). Herein they are jointly referred to as NMD-QTLs.

**Figure 1.**
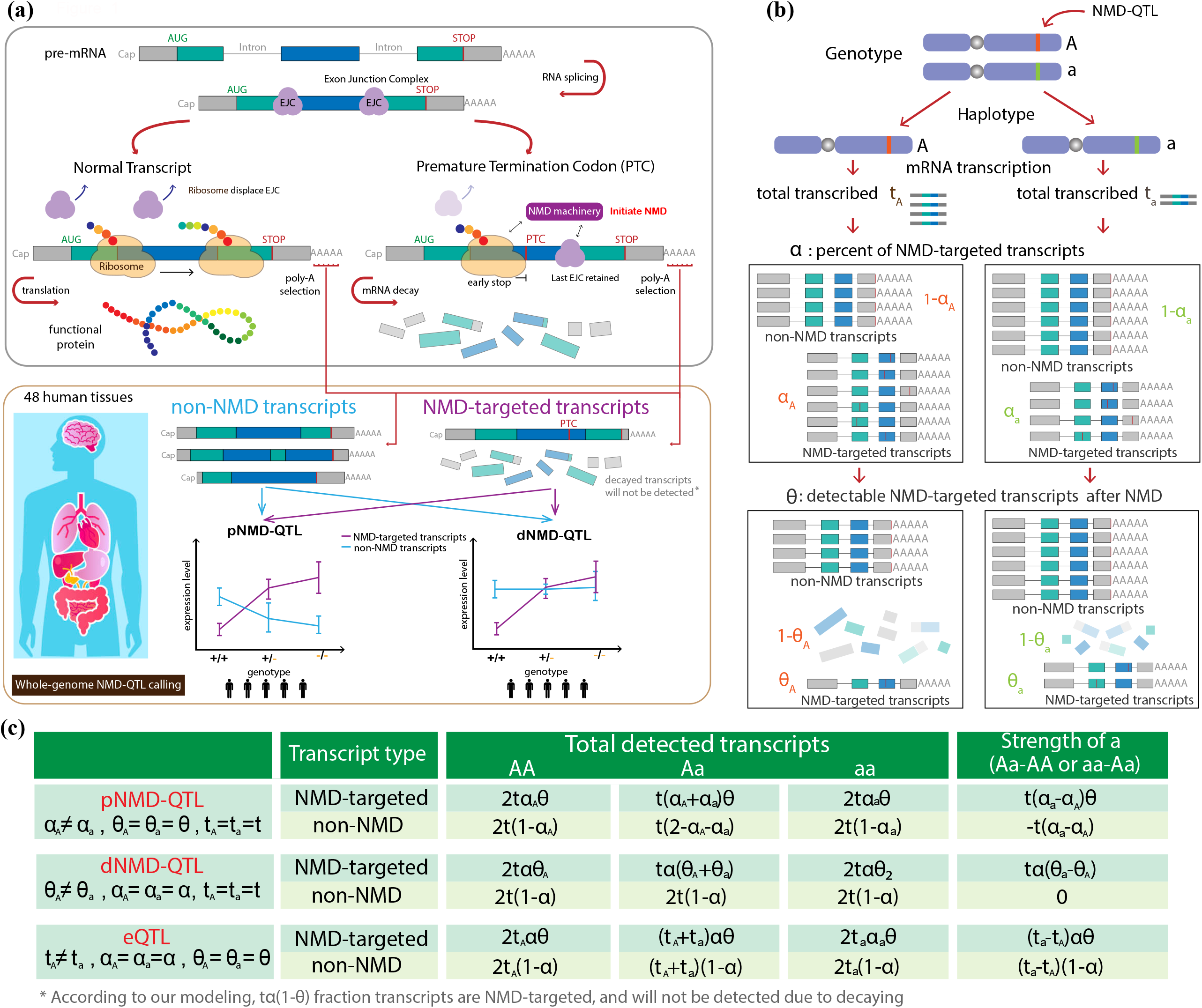
Modeling of NMD-QTLs. **(a)** Identification of NMD-QTLs across human tissues based on the GTEx data. **(b)** Parameterization of the expression of NMD genes. *t* represents the total amount of transcribed transcripts, *α* represents the percentage of transcribed transcripts targeted by NMD, and *θ* represents the percentage of NMD-targeted transcripts that remained after degradation. The parameters can be different for different alleles of a genetic variant. **(c)** Allelic effect sizes of a pNMD-QTL, dNMD-QTL, or eQTL for NMD-targeted transcripts and non-NMD transcripts.

As shown in Fig. 1c, pNMD-QTLs and dNMD-QTLs can be identified and distinguished from regular eQTLs by separately examining their allelic effect sizes for NMD-targeted transcripts and non-NMD transcripts. For a pNMD-QTL, the allelic effect sizes are in the opposite direction of NMD-targeted and non-NMD transcripts, and the observed regulatory strength is smaller for NMD-targeted transcripts as fewer are detected after degradation. For a dNMD-QTL, the significant association is only observable in NMD-targeted transcripts. Examples of a pNMD-QTL and a dNMD-QTL are shown in Fig. 2a. A positive effect size is observed for the alternative allele of the pNMD-QTL when considering NMD-targeted transcripts (i.e., higher transcript amount when an individual contains more copies of the alternative allele). However, the allelic effect size is negative for non-NMD transcripts. For the dNMD-QTL, the copy number of the alternative allele only affects the amounts of NMD-targeted transcripts.

**Figure 2.**
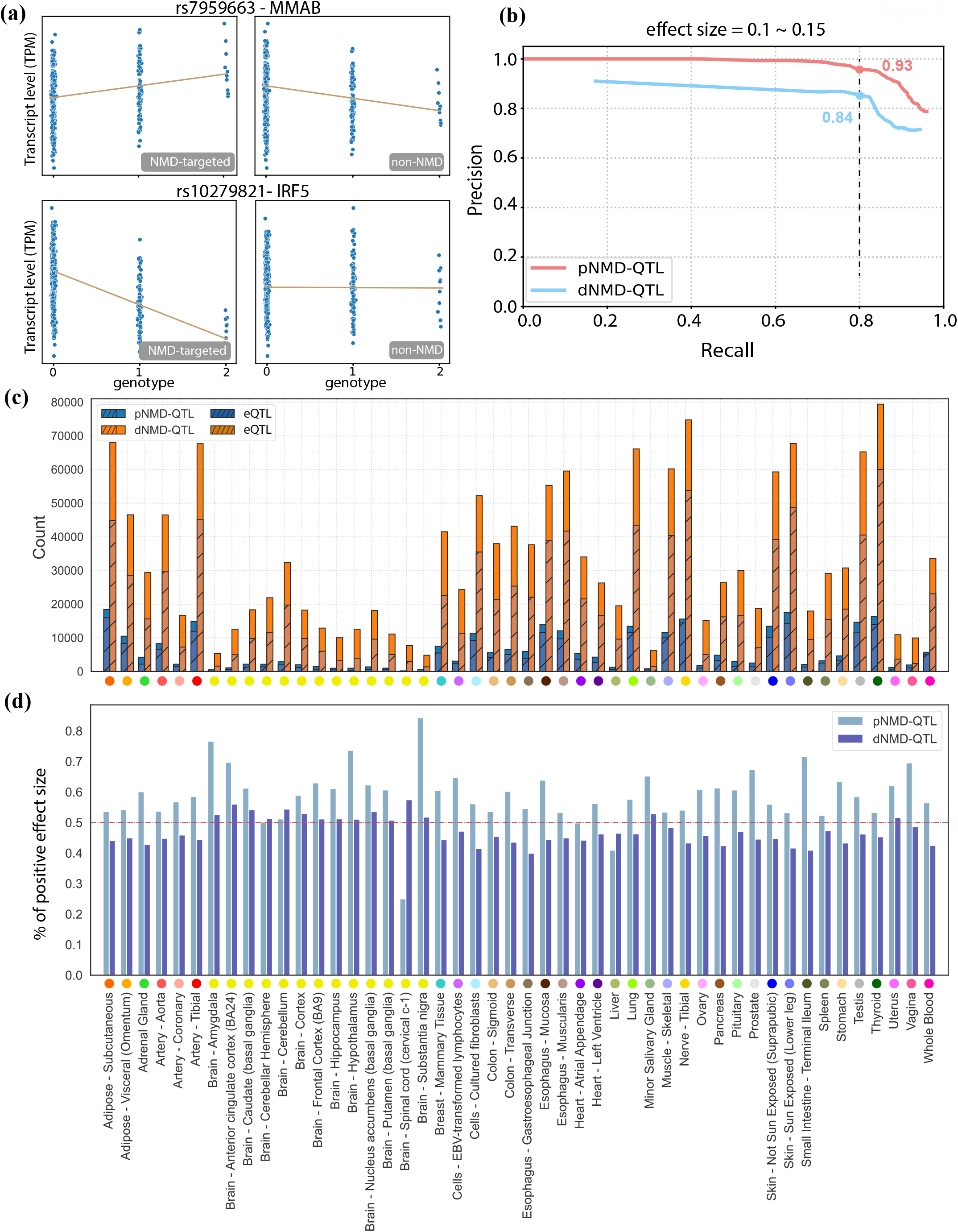
Identification of pNMD-QTLs and dNMD-QTLs. **(a)** Examples of a pNMD-QTL and a dNMD-QTL. A variant is identified as a pNMD-QTL if the allelic effect size is in opposite directions for NMD-targeted transcripts and non-NMD transcripts. A variant is identified as a dNMD-QTL if the association is only observed in NMD-targeted transcripts. **(b)** Precision-recall curves for identifying pNMD-QTLs and dNMD-QTLs. The effect size was simulated between 0.1 and 0.15. **(c)** The number of pNMD-QTLs and dNMD-QTLs detected for each of the 48 tissues. The shaded areas represent the number of variants also identified as eQTLs in GTEx. The color scheme used for tissues is the same as the one used in GTEx. **(d)** The fractions of positive effect sizes of pNMD-QTLs and dNMD-QTLs for NMD-targeted transcripts in each tissue.

### Evaluation of NMD-QTL Modelling

To evaluate the performance of our models, we estimated precision and recall for the identification of pNMD-QTLs and dNMD-QTLs from simulations. A total of 10,000 genetic variants were simulated; 5% were pNMD-QTLs (*α*_A_≠*α*_a_, *θ*_A_ =*θ*_a_, *t*_A_=*t*_a_), 5% were dNMD-QTLs (*α*_A_=*α*_a_, *θ*_A_ ≠*θ*_a_, *t*_A_=*t*_a_), and the remainder (90%) were null cases for NMD-QTLs (*α*_A_=*α*_a_, *θ*_A_ =*θ*_a_). The null cases were either regular eQTLs, or variants not associated with expression depending on the randomly assigned differences between *t*_*A*_ and *t*_*a*_. When the allelic effect size of NMD-QTLs was between 10% and 15% (i.e., |*α*_A_ − *α*_a_|=0.1-0.15 or |*θ*_A_ − *θ*_a_|=0.1-0.15), at a recall rate of 0.8, the precision was 0.93 and 0.84 for the identification of pNMD-QTLs and dNMD-QTLs, respectively (Fig. 2b). More detailed results and results for other simulations with smaller (5%-10%) or larger (15%-20%) allelic effect sizes can be found in Supplementary Fig. 1. The simulations verified we can distinguish NMD-QTLs from regular eQTLs by jointly considering the significance and directions of the allelic effect sizes for NMD-targeted and non-NMD transcripts. To ensure the accuracy of the effect size estimation, we applied an additional quantile regression model (see Methods) as quantile regression has the desired robustness to outliers and dropouts in transcriptomics data, and it significantly improves eQTL mapping with an accurate effect size estimation [38], which enables us to better identify NMD-QTLs.

### Identification of NMD-QTLs across different human tissues

We identified pNMD-QTLs and dNMD-QTLs in *cis* for each of the 48 tissues in GTEx v7 with the sample size ≥70. On average, 6033 pNMD-QTLs and 34049 dNMD-QTLs were detected for a tissue (Fig. 2c). Usually more dNMD-QTLs were discovered than pNMD-QTLs (4-11 folds in different tissues). The thyroid and tibial nerves had the highest numbers of NMD-QTLs (99053 and 95803 respectively), while the minor salivary gland and vagina tissues had the lowest (7044 and 12148 respectively). Worth noting is that GTEx, which models the overall expression as phenotypes without dissecting the NMD regulation, did not identify 7.3%-93.0% of pNMD-QTLs and 24.4%-76.0% of dNMD-QTLs as eQTLs (Fig. 2c, non-shaded areas).

To distinguish NMD-QTLs promoting or inhibiting NMD, we examined the effect size for NMD-targeted transcripts (Fig. 2d). Since the genotype was coded as the number of alternate non-reference alleles, a positive effect size suggests that *α*_*alt*_ > *α*_*ref*_ (pNMD-QTLs) or 1 − *θ*_*alt*_ <; 1 − *θ*_*ref*_ (dNMD-QTLs). An average of 59% pNMD-QTLs across tissues had positive signs. Thus, the alternate non-reference alleles promote the generation of NMD-targeted transcripts. Usually <50% of dNMD-QTLs had positive signs (average: 47%). Thus, the non-reference alleles of these dNMD-QTLs inhibit the decay of NMD-targeted transcripts. Equivalently, an average of 53% dNMD-QTLs had negative signs and their non-reference alleles promote the decay of NMD-targeted transcripts.

### Tissue specificity of NMD-QTLs

To examine the tissue similarity of NMD regulation, we considered a similarity score reflecting the enrichment of shared NMD-QTLs between two tissues (details in Methods). As shown in Fig. 3a, brain tissues had unique lists of NMD-QTLs: the NMD-QTL similarities were generally high among different brain tissues (average of 24.7), but the average similarity between brain and any other tissues was as low as 6.6. This suggests that NMD-QTLs were shared within different brain sub-regions but were substantially different from those in other tissues.

**Figure 3.**
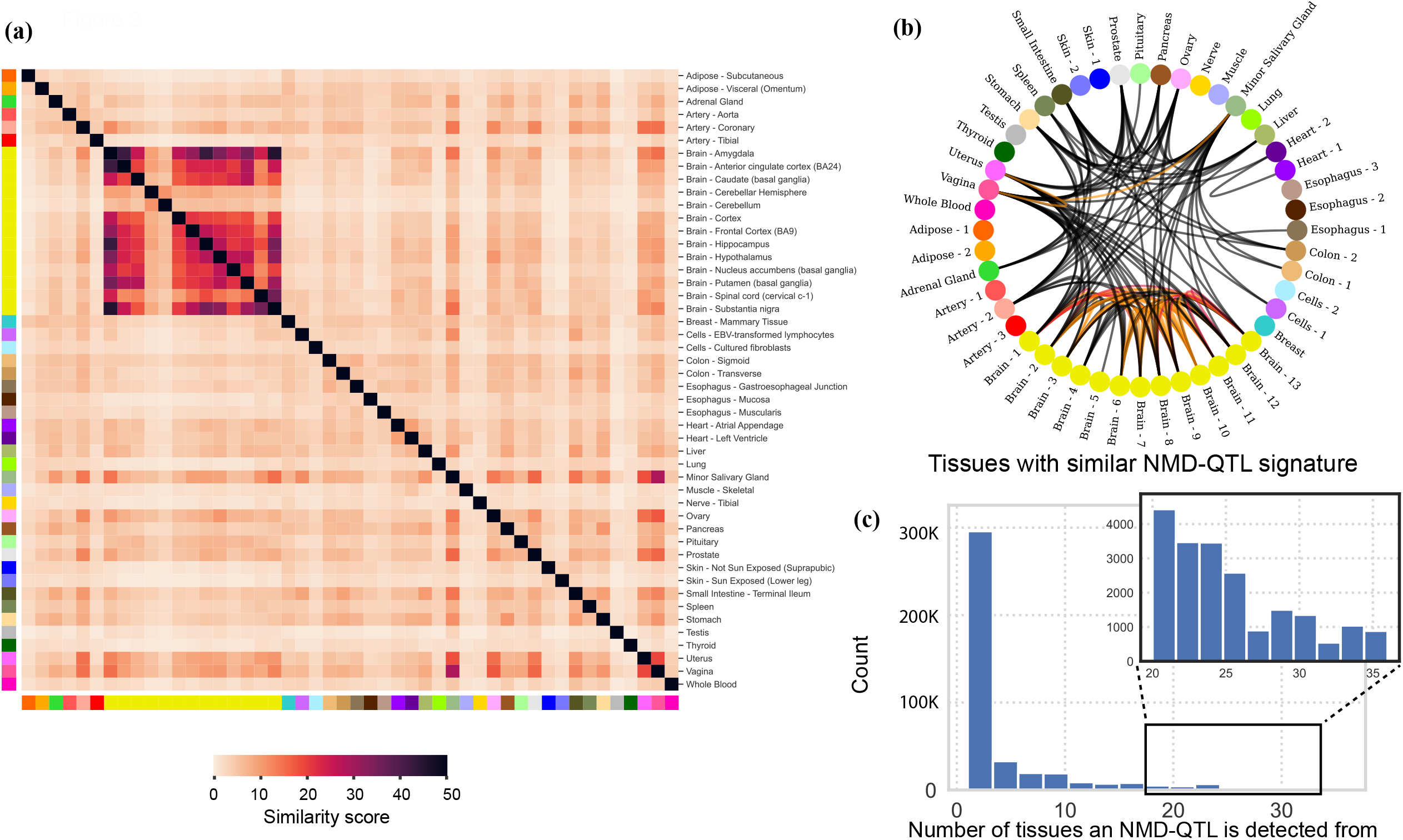
Tissue specificity of NMD-QTLs. **(a)** The similarity of NMD-QTL signatures between tissue pairs. The similarity score s* reflects the enrichment of shared NMD-QTLs among different tissues. The detected NMD-QTLs from brain tissues are very distinctive from other tissue. **(b)** Tissue interaction map. No edge is drawn if the similarity score s* between the two tissues <10. Black edges: 10 ≤ s* < 20; orange edges: 20 ≤ s* <30; and red edges: 30 ≤ s*. **(c)** The number of tissues in which an NMD-QTL variant is detected. Brain tissues are combined as one tissue.

The tissue interaction map based on the similarities of NMD-QTLs (Fig. 3b) clearly shows the distinction of brain tissues as well as the connections among major viscera (small intestine, spleen, stomach) and reproductive organs (uterus, vagina, ovary). The strong tissue specificity of NMD-QTLs is also supported by the observation that 54.3% of NMD-QTLs were discovered only in a single tissue (Fig. 3c, n.b. brain sub-regions were combined as one tissue). Taken together, NMD-QTLs exist as ubiquitously across the human body as NMD regulation itself, and many demonstrate strong tissue specificity.

A handful of NMD-QTLs were discovered in all tissues (Fig. 3c). These are located in important genes such as *TP53I13* (a tumor suppressor gene), *ST7L* (homologous to tumor suppressor gene *ST7*), *NDUFAF7* (an enzyme involved in the assembly and stabilization of Complex I, an important complex for mitochondrial respiratory chain), *IRF5* (belongs to a transcription factor family with diverse roles including virus-mediated activation of interferon, cell growth, differentiation, apoptosis, and immune system activity), *MMAB* (an enzyme that catalyzes the final step in the conversion of vitamin B_12_ into adenosylcobalamin), and three ribosomal proteins (S19, L13, L27a). It is known that NMD has high inter-individual variability and closely affects the clinical outcome for genetic disease [39,40]. Therefore, these natural variations of NMD-QTLs impacting tumor suppressors and important metabolic genes across all tissues may be exploited to enhance personalized cancer immunotherapy and to treat a wide range of genetic diseases.

### Colocalization of NMD-QTLs and disease SNPs

Functional variants are well-known to play an important role in diseases etiology. Utilizing the collection of disease-related SNPs in DisGeNET [41], we explored the colocalization of NMD-QTLs and disease-related SNPs. Specifically, out of the 54,643 non-redundant pNMD-QTLs and the 319,608 non-redundant dNMD-QTLs, 1151 (2.10%) and 3952 (1.24%) of them were found to be disease SNPs reported in DisGeNET, respectively (Fig. 4a). Compared to regular eQTLs discovered in GTEx for NMD-genes (0.82% were found to be disease SNPs), NMD-QTLs were much more likely to be colocalizing with disease SNPs: the odds ratio was 2.54 for pNMD-QLTs vs. eQTLs (p-value of the Fisher’s exact test: 8.5×10^−154^); and 1.49 for dNMD-QTLs vs. eQTLs (p-value of the Fisher’s exact test: 4.2×10^−94^).

**Figure 4.**
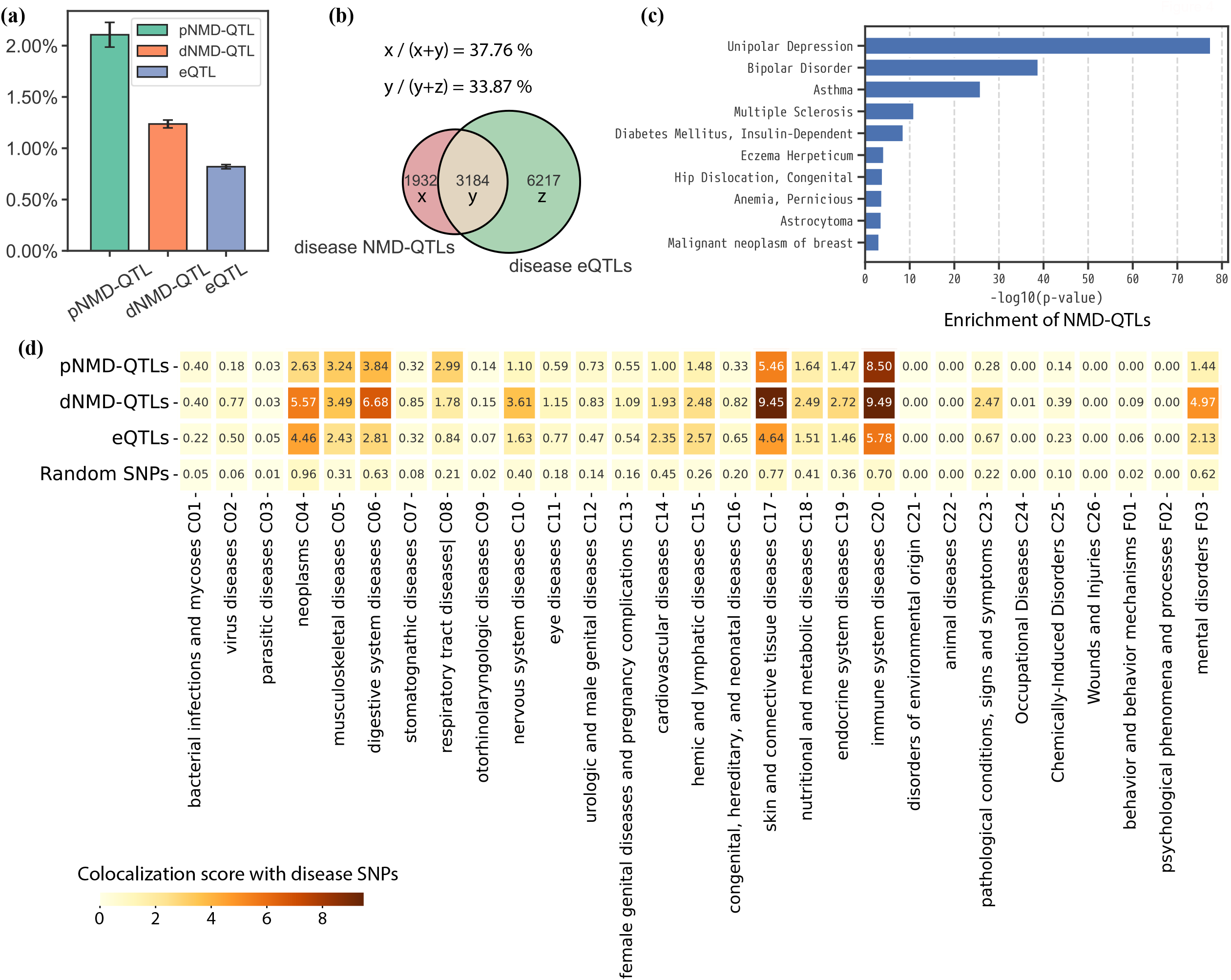
Variant-disease association analysis for NMD-QTLs. (**a**) Percentages of pNMD-QTLs, dNMD-QTLs, and eQTLs colocalizing with disease SNPs. (**b**) Venn diagram among disease-associated NMD-QTLs and disease-associated eQTLs. **(c)** Top 10 diseases associated with enriched NMD-QTLs. They are ranked by the p values of the proportion tests. **(d)** Colocalization with disease SNPs under each MeSH term. The colocalization scores are compared among pNMD-QTLs, dNMD-QTLs, eQTLs, and randomly sampled SNPs (the number is the same as that of NMD-QTLs).

Although the total number of non-redundant NMD-QTLs was only 11.53% of the total number of non-redundant eQTLs (331,233 vs. 2,872,430 for NMD-genes), we also discovered 33.87% disease SNPs colocalized with eQTLs by NMD-QTLs (Fig. 4b). More importantly, 37.76% disease SNPs colocalizing with NMD-QTLs were not detected as eQTLs (Fig. 4b). A total of 737 diseases were associated with NMD-QTLs, suggesting that NMD regulation is an essential component in maintaining the normal function of cells.

Among the 8073 total diseases considered in DisGeNET, we identified 43 with prominent NMD-QTLs enrichment (p-values < 0.05 for both the proportion test and the hypergeometric test, details in Supplementary Texts). Remarkably, 9 out of the 43 (20.9%) diseases were categorized as “mental disorders (F03)” under the “psychiatry and psychology (F)” category of the Medical Subject Headings (MeSH) system. The most significant was “unipolar depression”, followed by “bipolar disorder”. The top 10 diseases are shown in Fig. 4c and the complete list of 43 diseases is in Supplementary Table 1. Mutations in core NMD pathway genes are implicated in multiple mental disorders [42–45]. Together with our aforementioned findings that NMD-QTLs show special distinctiveness and tissue-specificity in brain tissues, the brain disease association further suggests the critical role of NMD regulation for brain function and malfunction.

The diseases in DisGeNet were categorized into 29 MeSH terms. We then studied the detailed colocalization status of disease SNPs and NMD-QTLs for each specific MeSH term. The calculation of the colocalization enrichment score is described in Methods. As shown in Fig. 4d, dNMD-QTLs were observed to be enriched in nervous system diseases (C10) and mental disorders (F03). The fold-change of the enrichment for dNMD-QTLs vs. eQTLs was 2.2 for C10 and 2.3 for F03. The fold change was as high as 9.0 for C10 and 8.0 for F03 when comparing dNMD-QTLs with randomly sampled SNPs. Contrariwise, pNMD-QTLs were less enriched than eQTLs for these two terms. But the fold change was still as high as 2.8 and 2.3, respectively, when compared to randomly selected SNPs. This suggests regulating the decay efficiency of NMD-targeted mRNA substrates is a key component of NMD regulation in the brain and neural systems. Additionally, dNMD-QTLs but not pNMD-QTLs were more enriched than eQTLs in terms such as neoplasm (C04) and pathological conditions (C23). Both pNMD-QTLs and dNMD-QTLs were more enriched than eQTLs for terms such as musculoskeletal diseases (C05), digestive system diseases (C06), respiratory tract diseases (C08), skin and connective tissue diseases (C17) and immune system diseases (C20).

Some NMD-QTLs were related to many diseases. For example, the pNMD-QTL SNP rs5443 was associated with 48 out of the total 8073 diseases. It acted as a pNMD-QTL to regulate the fraction of NMD-targeted transcripts of the DNA repair gene *XRCC3* in 18 tissues including 6 brain tissues. This implies that NMD regulation interplays closely with DNA repair and is essential in ensuring genome integrity and preventing genetic disease. The dNMD-QTL SNP rs1800629, regulating the degradation efficiency of NMD-transcripts of genes *HLA-C* and *ATP6V1G2-DDX39B* (read-through transcription between neighboring genes *ATP6V1G2* and *DDX39B*), was related to 58 diseases, 18 of which were related to neoplastic processes.

### Disease-associated NMD-QTLs in brain tissues

Compared to non-disease NMD-QTLs, disease-associated NMD-QTLs showed an even stronger tissue specificity in brain sub-regions (Fig. 5a). By contrast, brain compartmentalization was weaker in disease eQTLs than in non-disease eQTLs. This reaffirms that NMD-QTLs play a critical role in maintaining neural system functionalities and can be valuable for examining brain and psychiatric pathogenesis.

**Figure 5.**
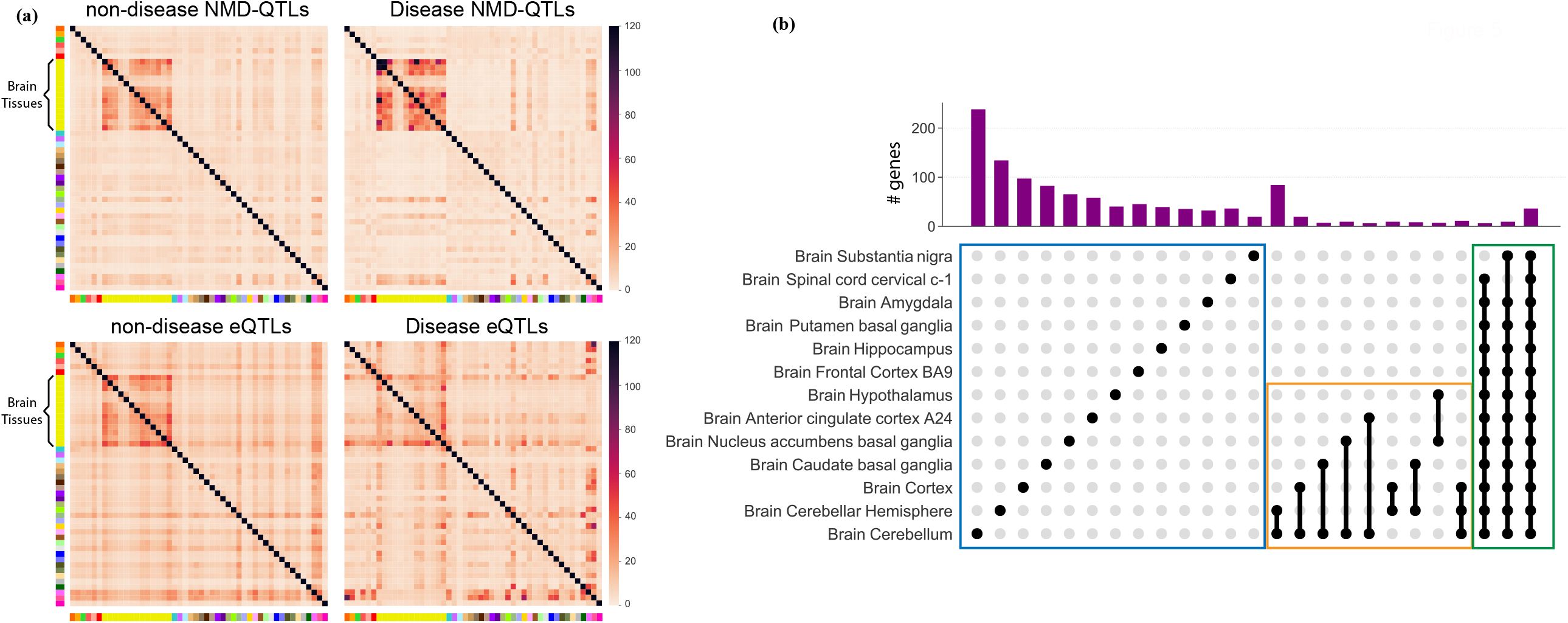
Disease NMD-QTLs in brain tissues. **(a)** Tissue pair-wise similarity of non-disease and disease QTL signatures. Disease NMD-QTLs but not disease eQTLs are more similar in brain tissues. **(b)** UpSet plot showing the top 25 most frequent brain sub-region memberships displayed by genes with NMD-QTLs.

We examined the sub-region membership of genes with at least one discovered NMD-QTL (i.e., the specific set of brain sub-regions from which NMD-QTLs were discovered for a given gene). The 25 most frequent brain sub-region memberships were displayed via an UpSet plot (Fig. 5b). The most frequent membership pattern appeared to be genes with NMD-QTLs identified in a single or a few sub-regions (blue and orange boxes in Fig. 5b), as well as genes with NMD-QTLs discovered in most brain sub-regions (green box in Fig. 5b).

### NMD-QTLs are more likely to be located within gene bodies and exons compared to eQTLs

The 50-nt rule was applied to tag NMD-targeted transcripts in GENCODE because the rule has the strongest predictive value for NMD susceptibility [46]. In this study, NMD-targeted transcripts all follow this rule but the percentage of transcripts undergoing NMD and their decay efficiency can vary. The genomic positions of NMD-QTLs can inform additional determinants of NMD regulation.

First, we examined whether NMD-QTLs had any preference for residing inside a gene body or in the intergenic regions. For each gene, we counted the numbers of NMD-QTLs located within (N_i,g_) or outside (N_o,g_) of the gene body. Then a linear model was fit between N_i,g_ and N_o,g_ taking the gene length (l_g_) into account: *N*_*i,g*_ = *w*_0_ + *w*_1_*N*_0,*g*_ + *w*_2_*log*_2_(l_*g*_). We modeled each tissue separately and compared the results for NMD-QTLs and eQTLs. As shown in Fig. 6a, the estimations of *ŵ*_1_ for NMD-QTLs were consistently larger than those for eQTLs, especially for brain tissues (Fig. 6c). This suggests that, compared to eQTLs, NMD-QTLs are more likely to function within a gene body than in intergenic regions. However, the estimations of *ŵ*_2_ for NMD-QTLs and eQTLs were similar (p-value=0.37, Wilcoxon test, Fig. 6b). This suggests that the gene length factor was controlled at a similar level for both.

**Figure 6.**
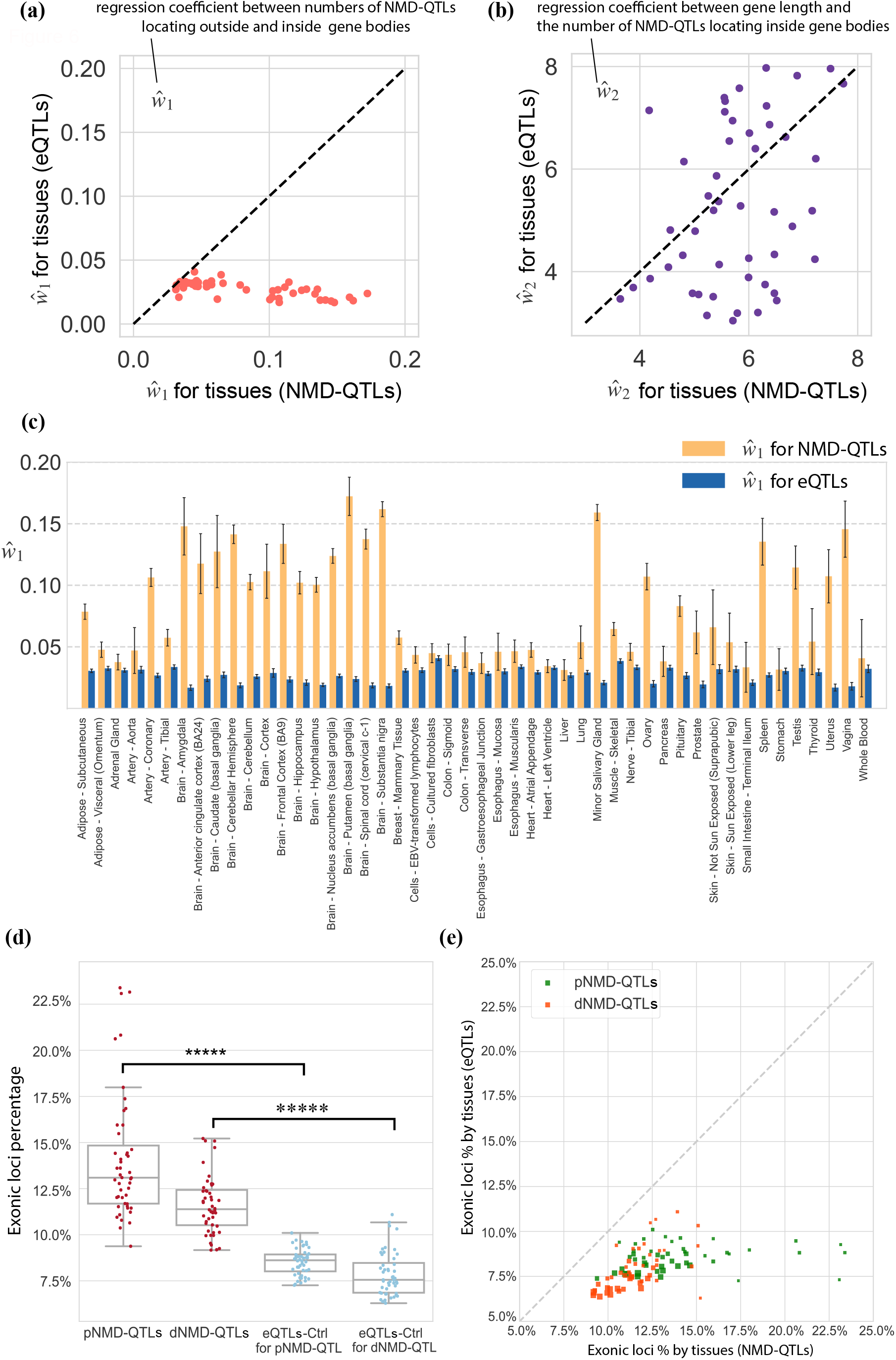
Genomic locations of NMD-QTLs. **(a)** The coefficient estimations for 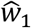. **(b)** The coefficient estimations for 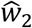. 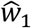 and 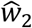 are for the model *N*_*i,g*_ = *w*_0_ + *w*_1_*N*_0,*g*_ + *w*_2_lo*g*_2_(l_*g*_), where N_i,g_ is the number of NMD-QTLs (or eQTLs) inside a gene, N_o,g_ is the number of NMD-QTLs (or eQTLs) outside of any gene, and l_*g*_ is the gene length. **(c)** The 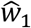 estimation for each tissue. Error bars: 95% confidence intervals. The 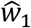 estimations for NMD-QTLs are consistently larger than those for eQTLs, meaning that NMD-QTLs prefer locating inside a gene body. **(d)** The proportion of genic variants that are located on exons. The exonic proportions are significantly higher for NMD-QTLs, compared to eQTLs discovered for the same genes. **(e)** The scatter plots of the exonic proportions. The size of the symbol is proportional to the sample size for a specific tissue.

For those NMD-QTLs residing within a gene body, we further examined whether they prefer exon or intron regions. Compared to eQTLs, NMD-QTLs were more likely to be in exonic positions (Fig. 6d, p-values from Wilcox tests <1×10^−6^). The tissue-wise exonic proportions of NMD-QTLs and eQTLs are compared in Fig. 6e, further supporting the exonic location preference of NMD-QTLs. Regulatory elements for mRNA degeneration are usually exonic after mRNA splicing. In addition, exon-junction complex (EJC) has been reported to be involved in the initiation of NMD [47,48]. It is possible that NMD-QTLs in exonic regions interact more effectively with the deposition of EJC to tag NMD regulation for natural transcripts.

### Ordinal positions of NMD-QTLs along exons and introns

Since the location of EJC is essential for NMD regulation, we next studied the relative positions of NMD-QTLs within a gene body by defining an ordinal rank for exons and introns. For each gene, the combined exon intervals from the collapsed gene model (i.e., combining all isoforms of a single gene into a single transcript) used in GTEx were considered. The intron intervals are deduced from those exon intervals. We ranked these exon and intron intervals separately from both 5’-end and 3’-end, starting from 0. For all genes with NMD-QTLs, the frequency count and the median length of the exon (or intron) intervals on each ordinal position are shown in Supplementary Figs 2a,b. We recorded exon (or intron) ordinal positions for all genic NMD-QTLs; the raw count distributions are shown in Supplementary Figs 2c,d.

To remove counting bias, we normalized the raw count by the number of exon or intron intervals at each ordinal position (Fig. 7a,b), and further by the median length of exon or intron intervals at each ordinal position (Fig. 7c,d). Both pNMD-QTLs and dNMD-QTLs were more likely to be located in the last exon interval (rank 0 at 3’, Fig. 7a). This was partly due to the fact that the last exon interval is usually much longer than other exons (median length of 831 vs. 187). Taking the exon interval length into account, the chance of an NMD-QTL locating in an exonic position of the last exon interval (rank 0 at 3’) was actually much lower (Fig. 7c). The highest frequency was within the penultimate exon from the 3’ end (i.e., rank 1), followed by the two upstream exons (i.e., ranks 1-2). These 3’ exon ordinal positions may influence the formation of the last EJC or affect the interaction between PTC and downstream EJC. As regards the intron ordinal positions for NMD-QTLs, both pNMD-QTLs and dNMD-QTLs preferred the first intron interval (rank 0 at 5’, Fig 7b,d).

**Figure 7.**
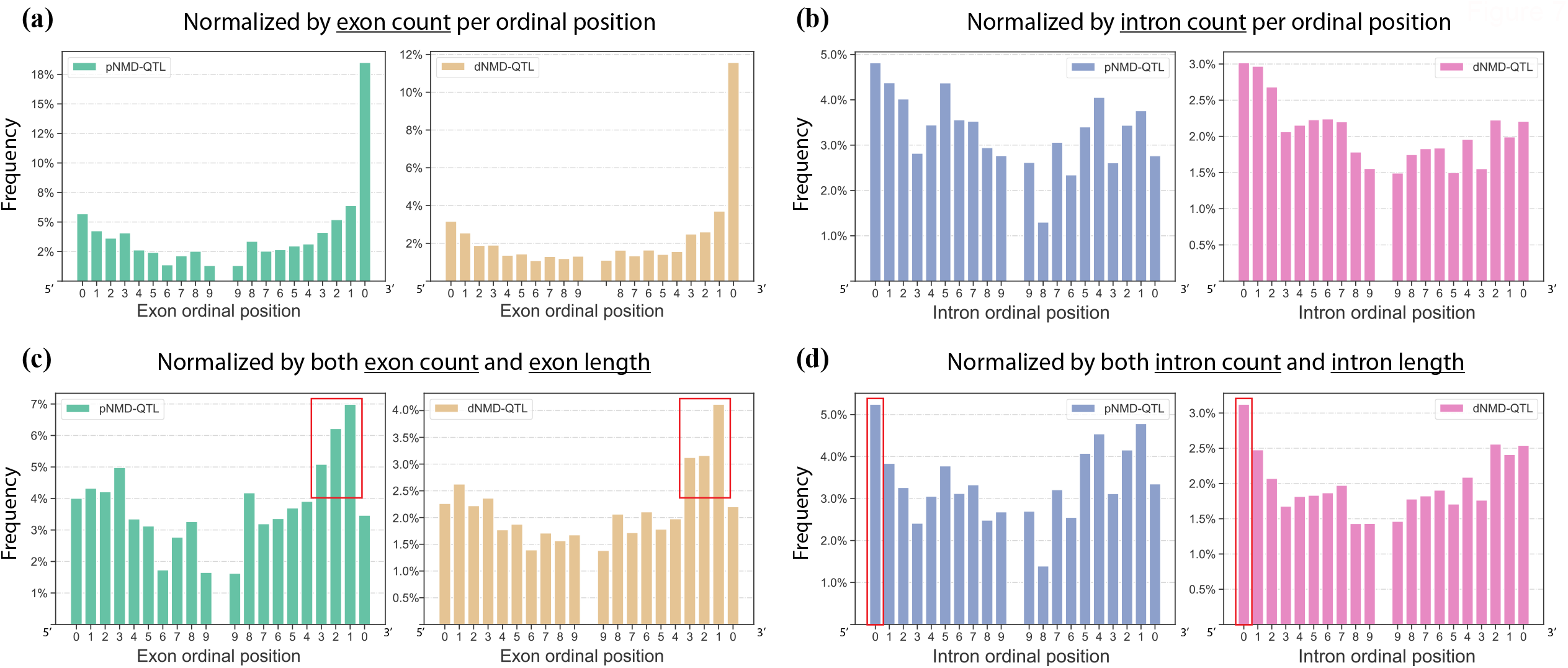
Ordinal positioning of NMD-QTLs along exon and intron intervals. The exon and intron intervals are deduced from the collapsed gene model and ranked from 0 for both 5’ and 3’ ends. **(a)** Distribution of the count of NMD-QTLs on each exon ordinal position normalized by the exon count per ordinal position. **(b)** Distribution of the count of NMD-QTLs on each intron ordinal position normalized by the intron count per ordinal position. **(c)** Distribution of NMD-QTL counts on each exon ordinal position normalized by both the exon count and exon length per ordinal position. **(d)** Distribution of NMD-QTL counts on each intron ordinal position normalized by both the intron count and intron length per ordinal position.

### Location preference of NMD-QTLs at the base-pair (bp) resolution

In addition to the ordinal positions along exons or introns, we examined the base-pair (bp) distance of NMD-QTLs to the exon or intron boundary. Both exonic NMD-QTLs and eQTLs favored the 0-50 bp regions close to exon boundaries (Fig. 8a for pNMD-QTLs and Supplementary Fig. 3a for dNMD-QTLs). On the other hand, intronic NMD-QTLs were more likely to be near intron boundaries than intronic eQTLs (Fig. 8b and Supplementary Fig. 3b).

**Figure 8.**
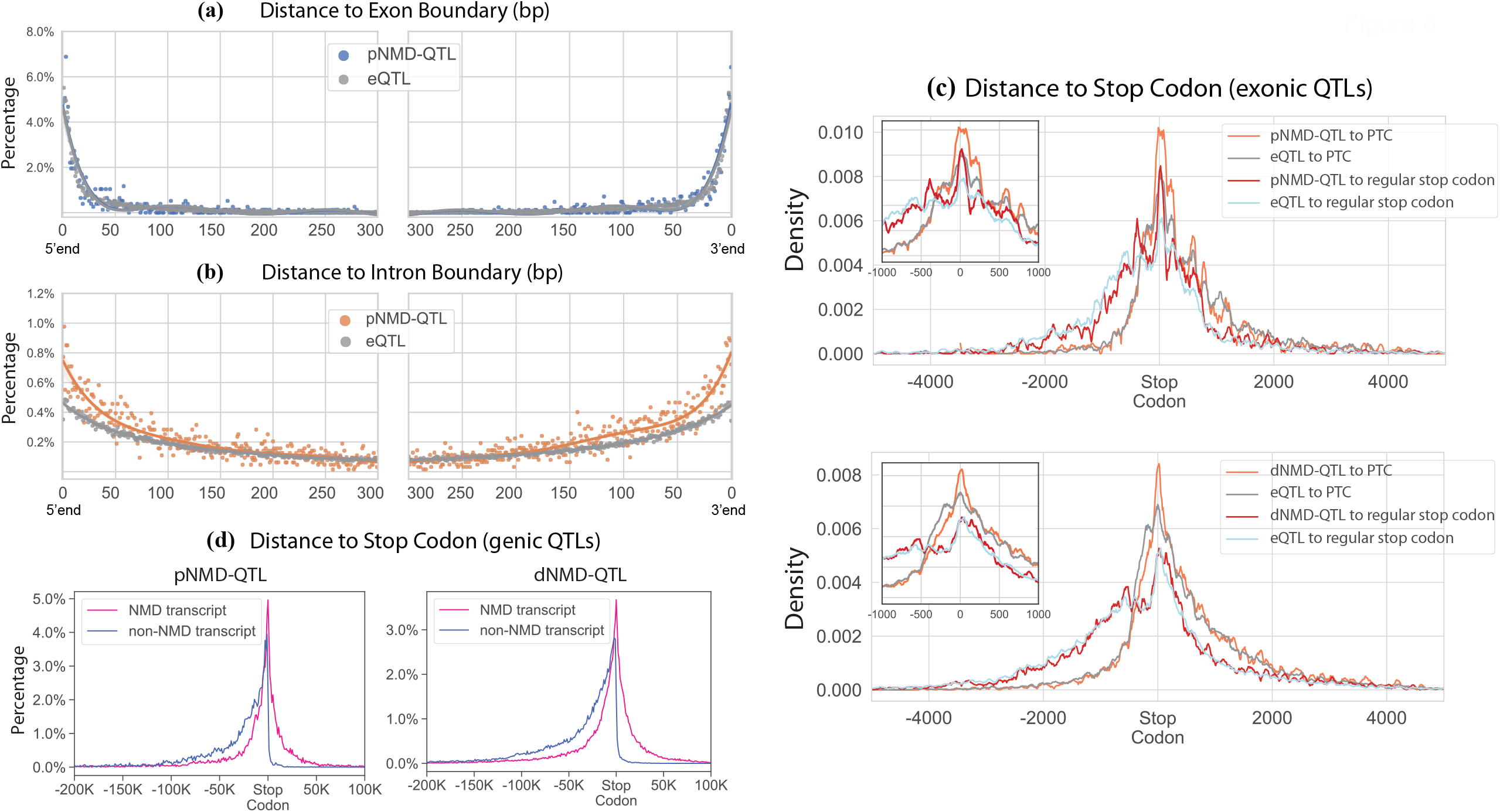
Distances of NMD-QTLs to boundaries of exons, introns, and stop codons. **(a)** Distances to exon boundaries for variants located on an exon. The comparison is made between pNMD-QTLs and eQTLs. **(b)** Distances to intron boundaries for variants located on an intron. The comparison is made between pNMD-QTLs and eQTLs. Similar results for dNMD-QTLs can be found in Supplementary Figure 3. **(c)** Distances of exonic NMD-QTLs to PTCs of NMD-targeted transcripts, and to regular stop codons of non-NMD transcripts. Only exonic positions are counted in the distance calculation. **(d)** Distance between genic NMD-QTLs and the stop codons. Both exonic and intronic positions are counted in the distance calculation.

The location of PTC is tightly related to NMD regulation. We therefore considered the distances between NMD-QTLs and PTCs of NMD-targeted transcripts (vs. regular stop codons of non-NMD transcripts). For exonic NMD-QTLs, we calculated the distances on the transcripts without considering intronic positions (Fig. 8c). Compared to eQTLs, pNMD-QTLs were more concentrated in the (−75 bp, +150 bp) regions around PTCs (orange line vs. gray line in the upper panel of Fig. 8c). Interestingly, pNMD-QTLs were also more concentrated around the (−25 bp, +50 bp) regions around regular stop codons than eQTLs (red line vs. light blue line). We also observed another pNMD-QTLs peak (red line) around 400 bp upstream of the regular stop codons. Since the median distance between the upstream PTC and the downstream regular stop codon of the same gene is 822 bp (Supplementary Fig. 4), the peak on −400 bp of regular stop codons usually corresponds to positions downstream of PTCs.

The exonic dNMD-QTLs were only slightly more concentrated around PTCs (orange line vs. gray line in the lower panel of Fig. 8c). Their distribution is almost indistinguishable around regular stop codons (red line vs. light blue line in the lower panel of Fig. 8c). We also considered the genomic distances (both exonic positions and intronic positions) between genic NMD-QTLs and PTCs (or regular stop codons). As shown in Fig. 8d, NMD-QTLs can be located upstream or downstream of PTCs but are rarely downstream of regular stop codons.

### The relationship between NMD-QTLs and miRNAs or RNA-binding proteins (RBPs)

We wonder how these *cis*-acting NMD-QTLs interact with *trans*-acting factors to impact NMD regulation. We focused on miRNAs and RBPs. miRNAs recognize complementary mRNA molecules and function in RNA silence and gene regulation [49,50]. RBPs are critical for post-transcriptional RNA processing and modification (e.g., splicing, mRNA stabilization and mRNA localization) [51,52]. We hypothesize some NMD-QTLs could function by affecting the miRNA or RBP binding sites. Here we examined how many NMD-QTLs were located at miRNA target regions predicted by TargetScan [53] as well as how many NMD-QTLs were located at RBP binding regions surveyed in ENCODE [54].

When we considered the targets of conservative miRNAs, 7.6‰ nonredundant pNMD-QTLs and 6.3‰ nonredundant dNMD-QTLs were found to be located in the miRNA target regions. Both were higher than the 4.2‰ level observed for regular eQTLs (for pNMD-QTLs vs. eQTLs, odds ratio=1.79, Fisher’s exact test p-value=6.3×10^−29^, for dNMD-QTLs vs. eQTLs, odds ratio=1.49, Fisher’s exact test p-value=1.5×10^−62^). The top 20 miRNAs whose targets were enriched with pNMD-QTLs or dNMD-QTLs are shown in Fig. 9a. Seven of them were shared between pNMD-QTLs and dNMD-QTLs (orange colored). The full list of miRNAs can be found in Supplementary Table 2. When the colocalization percentages were calculated for individual tissues, NMD-QTLs still had higher percentages than eQTLs (Fig. 9b).

**Figure 9.**
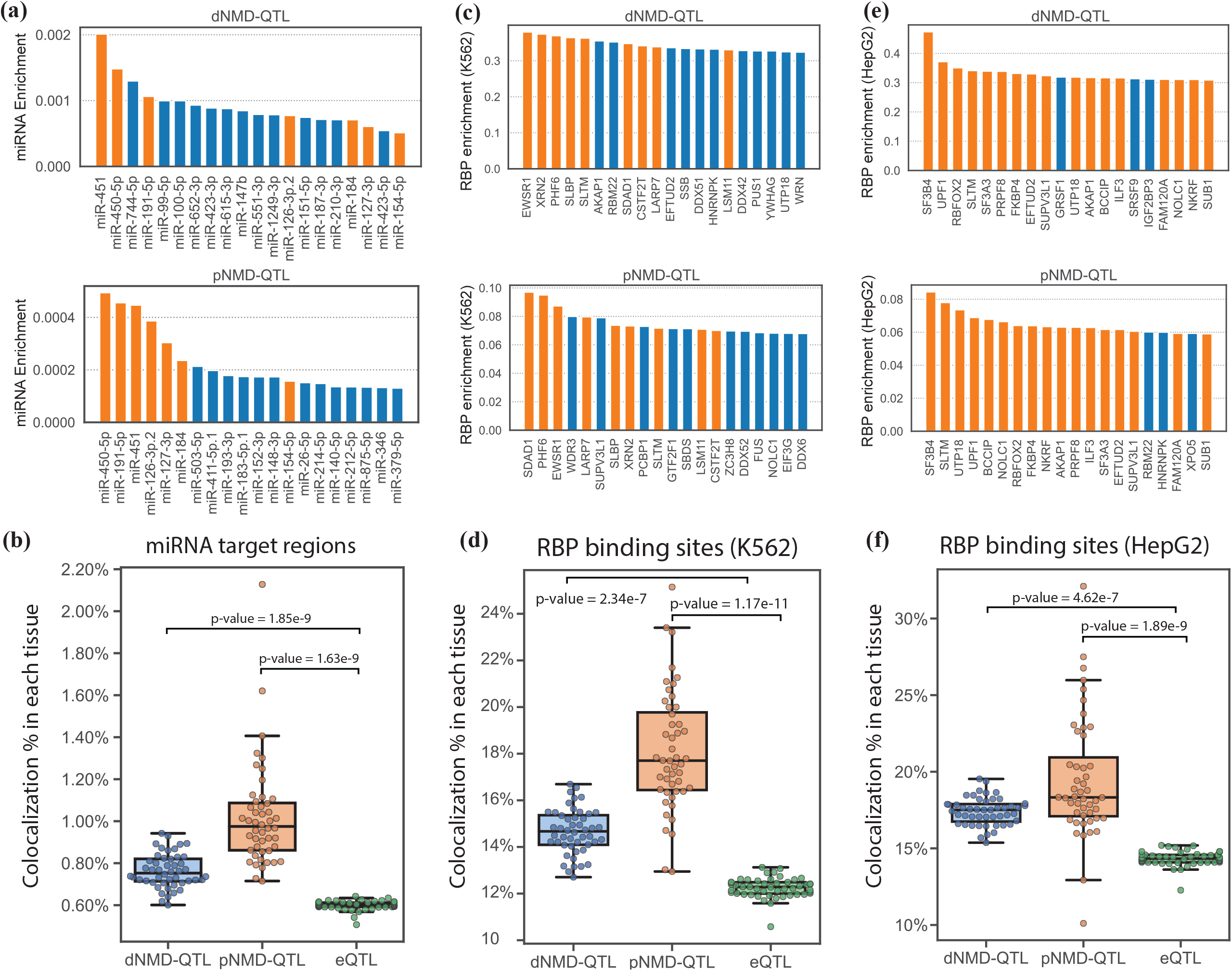
Relationship between NMD-QTLs and miRNAs or RBPs. **(a)** Top 20 miRNAs whose target regions are enriched with dNMD-QTLs or pNMD-QTLs. Shared miRNAs between dNMD-QTLs and pNMD-QTLs are orange colored. **(b)** Percentages of NMD-QTLs in miRNA targets compared to that of eQTLs. **(c)** Top 20 RBPs whose binding regions profiled in K562 are enriched with dNMD-QTLs or pNMD-QTLs. **(d)** Percentages of NMD-QTLs in binding regions of RBPs in K562 compared to that of eQTLs. **(e**) Top 20 RBPs whose binding regions profiled in HepG2 are enriched with dNMD-QTLs or pNMD-QTLs. (**d**) Percentages of NMD-QTLs in binding regions of RBPs in HepG2 compared to that of eQTLs.

A total of 12.7% nonredundant dNMD-QTLs and 14.9% nonredundant pNMD-QTLs were found in the binding sites of RBPs in the K562 cell line. Similarly, 14.9% nonredundant dNMD-QTLs and 16.4% nonredundant pNMD-QTLs were found in the binding sites of RBPs in the HepG2 cell line. These percentages were higher than those of eQTLs (10.3% and 12.3% in K562 and HepG2, respectively). For pNMD-QTLs vs. eQTLs, the odds ratio was 2.84 for K562 and 3.13 for HepG2 (Fisher’s exact test p-values <10^−16^). For dNMD-QTLs vs. eQTLs, the odds ratio was 2.42 for K562 and 2.84 for HepG2 (Fisher’s exact test p-values <10^−16^). Nine of the top 20 RBPs whose targets in the K562 cell line were enriched with pNMD-QTLs or dNMD-QTLs were shared (Fig. 9c, orange colored). The top 20 RBPs in HepG2 were similar between pNMD-QTLs and dNMD-QTLs (17 out of 20 were shared and orange colored, Fig. 9e), including UPF1, the key effector of the NMD pathway. Other RBPs could function as co-factors or modulators of the NMD regulation pathways. Full lists of RBPs are provided in Supplementary Table 3. When we considered NMD-QTLs discovered from each individual tissue instead of pooling them together, we still found that NMD-QTLs, especially pNMD-QTLs, were more likely to be in the binding sites of RBPs compared to eQTLs (Figs. 9d,f).

### Colocalization of splicing-QTLs and NMD-QTLs

In mammalian cells, alternative splicing coupled to nonsense-mediated decay (i.e., AS-NMD) is a conserved post-transcriptional regulation mechanism in which alternative splicing can alter the reading frame to produce a PTC-containing transcript subject to NMD [24,55–59]. We therefore analyzed the colocalization of NMD-QTLs and splicing-QTLs. Here, we identified splicing-QTLs by mapping variants associated with the inclusion ratios of annotated cassette exons (details in Methods). As expected, NMD-QTLs were much more likely to be colocalized with splicing-QTLs than with eQTLs (average odds ratios: 15.9 for pNMD-QTLs vs. eQTLs; 6.1 for dNMD-QTLs vs. eQTLs). Compared to dNMD-QTLs, pNMD-QTLs were more likely to act as splicing-QTLs simultaneously since alternative splicing is coupled with NMD through ‘producing’ transcripts subject to NMD. Such co-localization of pNMD-QTLs and splicing-QTLs was more prominent in brain tissues (average odds ratio in brain tissues for pNMD-QTL vs. eQTLs: 30.8, Fig. 10a). The odds ratio was as high as 60.8 in the amygdala which defines and regulates human emotions. The results suggest that AS-NMD can be a prevalent post-transcriptional regulation mechanism for brain tissues, especially in the amygdala.

**Figure 10.**
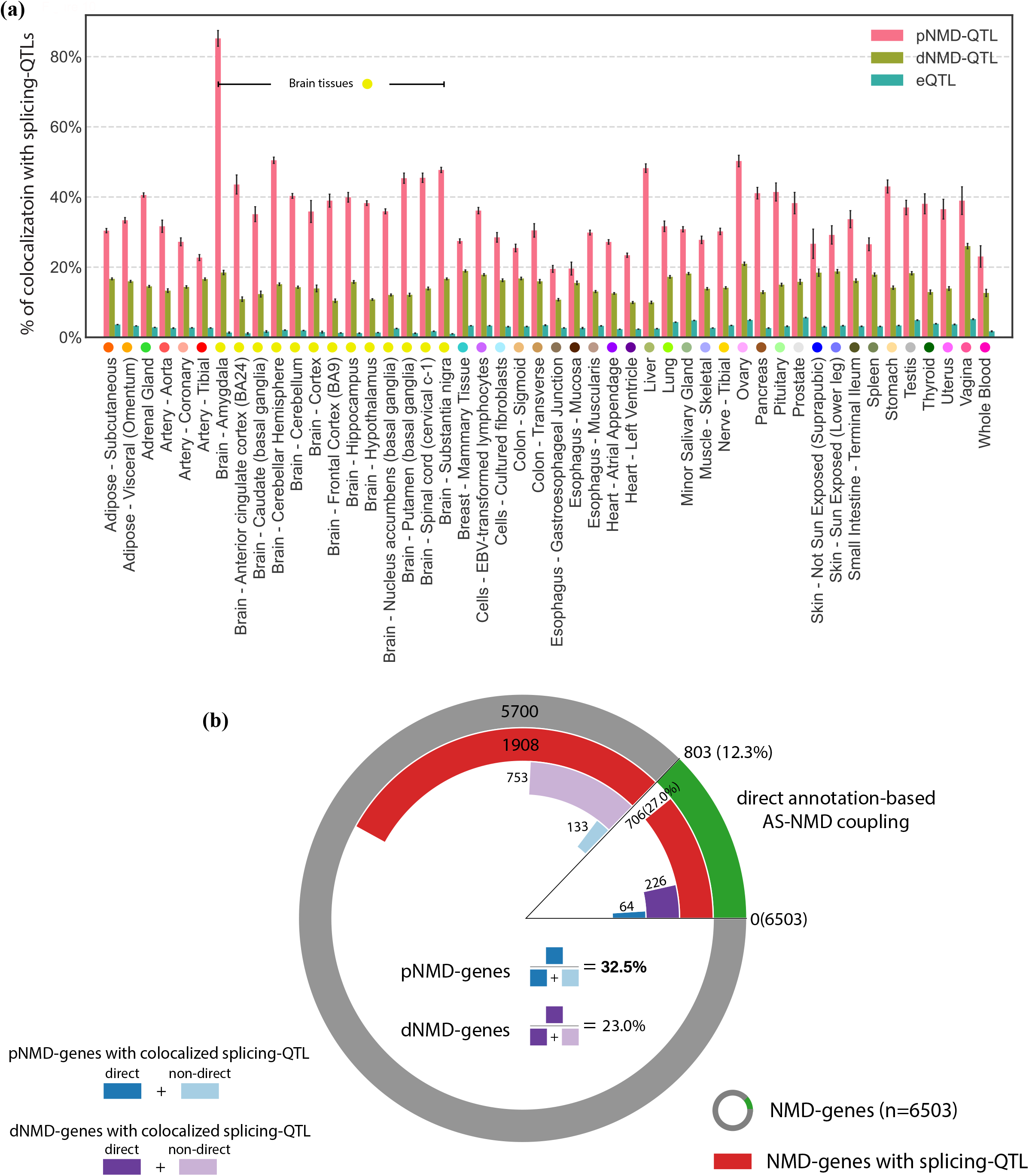
Colocalization of NMD-QTLs and splicing-QTLs. **(a)** Percentages of NMD-QTLs and eQTLs colocalizing with splicing-QTLs across tissues. Error bars: 95% confidence intervals. **(b)** Nested pie chart showing the relative proportion of each category. The most outer circle shows NMD-genes with and without annotation-based AS-NMD coupling (i.e., direct or non-direct). The inner circles from outside to inside shows NMD-genes with splicing-QTLs (red arcs), dNMD-genes (i.e., genes with dNMD-QTLs) with colocalized splicing-QTLs (purple arcs), and pNMD-genes (i.e., genes with pNMD-QTLs) with colocalized splicing-QTLs respectively (blue arcs).

To validate this finding, we focused on NMD-genes with direct annotation-based evidence of AS-NMD coupling: given a cassette exon, all NMD-targeted transcripts include the cassette exon and all non-NMD transcripts skip that cassette exon, or conversely, all NMD transcripts skip the cassette exon and all non-NMD transcripts include it. As shown in Fig. 10b, out of all 6503 NMD-genes, 803 (12.3%) of them had direct annotation evidence for AS-NMD coupling (the green arc). Out of the 2614 NMD-genes with splicing-QTLs (the red arcs), 706 genes (27.0%) showed direct evidence of AS-NMD coupling. For genes regulated by colocalized pNMD-QTLs and splicing-QTLs (the blue arcs), the percentage of genes with direct AS-NMD coupling was even higher (64 out of 197, 32.5%). However, this percentage was only 23.1% (226 out of 979) for genes with colocalized dNMD-QTLs and splicing-QTLs (the purple arcs). Therefore, NMD-genes with direct AS-NMD coupling were more likely to have splicing-QTLs and colocalized pNMD-QTLs (Fisher’s exact tests p-values <10^−16^). This corroborates our hypothesis that alternative splicing couples with NMD through ‘producing’ NMD-transcripts and suggests that alternative splicing itself may not influence decay efficiency of NMD transcripts.

## Discussion

To uncover the etiological path from SNPs to resultant phenotypic traits, it is essential to understand the mechanisms underlying gene expression variation. The current gene expression regulatory landscape derived from eQTL studies is far from complete. Traditional eQTL studies have focused on the identification of genetic loci linked to variations in the overall mRNA expression [60–62], and more recently to variations in mRNA splicing [63– 65]. A special interest of the current human eQTL studies is tissue-specific genetic impacts such as the efforts in GTEx [60,66,67], because they can provide valuable insights into disease phenotypes manifested on particular tissues. However, transcriptional and post-transcriptional regulations jointly determine tissue-specific gene expression, which needs a more careful integrated examination. In this study, we analyzed the post-transcriptional processing of RNA and proposed a more fine-grind regulatory QTL model, the NMD-QTL model, to pinpoint genetic variants that function via regulating the production percentage of NMD-transcripts (pNMD-QTLs) and the decay efficiency of targeted RNAs (dNMD-QTLs). Through joint consideration of allelic effect sizes for both NMD-transcripts and normal transcripts, we distinguish pNMD-QTLs and dNMD-QTLs from regular eQTLs. The modelling is validated by extensive simulations mimicking real data sets. Taking advantage of the unprecedented resource available in GTEx, we reveal the genome-wide landscape of genetic variants affecting NMD regulation which can be missed in regular GTEx eQTLs mapping, and uncover its tissue specificity and its variant-disease associations.

Our results show NMD-QTLs are valuable in dissecting the variant-disease associations and to characterize disease susceptibility, especially for brain and mental disorders. The higher colocalization rates of disease SNPs with NMD-QTLs than with eQTLs point to the need to include NMD-QTLs in prioritizing and interpreting variants discovered from the genome-wide association studies. The enrichment of NMD-QTLs was observed for a variety of diseases, including musculoskeletal diseases, digestive system diseases, respiratory tract diseases, skin and connective tissue diseases, and immune system diseases. The link between NMD regulation and various disease susceptibility stresses the fact that NMD functions as an intricate gene regulation mechanism across human tissues. In particular, brain tissues exhibit unique NMD-QTL signatures, and these NMD-QTLs can uncover prominent variant-disease associations.

Approximately 3 million individuals in the US alone are afflicted with genetic diseases caused by nonsense mutations (PTC diseases) [68]. Additionally, deficiencies in NMD factors are linked to neurological disorders, immune diseases, and cancers [15,69,70]. An NMD-based therapeutic strategy will be impossible without a comprehensive understanding of NMD regulation. First, NMD efficiency influences patients’ response to nonsense-suppression drugs [71], but NMD efficiency modulation is unclear. Second, NMD inhibitors have been proposed to treat PTC diseases, but potential side effects are awaiting to be addressed [29,68,72]. Our discovered NMD-QTLs may assist in the design of gene-specific NMD inhibition to treat PTC diseases, mitigating the global side effects caused by manipulations of core NMD factors [73].

Little is known about the determinants of NMD efficiency. The enrichment of NMD-QTLs within gene bodies and exons, especially the penultimate exons from the 3’ end, suggests embedded sequence features for NMD regulation. Regions around PTC are important regulatory regions since NMD-QTLs tend to be located around PTCs. The enrichment of pNMD-QTLs in near proximity to regular stop codons and their upstream 400 bp positions implies additional determinants producing NMD-targeted transcripts.

The location preferences of NMD-QTLs in the targets of miRNAs and RBPs show that NMD regulation is not fixated but can be modulated by miRNAs and RBPs. The mammalian core NMD machinery includes a PIKK complex (SMG1, SMG8, SMG9) and other SMG proteins (SMG5, SMG6, SMG7), UPF proteins (UPF1, UPF2, UPF3A, UPF3B), eukaryotic release factors (eRF1, eRF3), and exon junction complex (EJC) members (eIF4A3, RBM8A, MAGOH, and MLN51) [13,61,62]. Enrichments of NMD-QTLs colocalizing with RBP binding sites and miRNA targets suggest that these *cis*-acting regulatory elements interact with RBPs and miRNAs to modulate the generation and decay efficiency of NMD transcripts. Additionally, the colocalization of NMD-QTLs especially pNMD-QTLs with splicing-QTLs suggests that RBPs may interact with NMD regulations through the coupling of alternative splicing and NMD.

## Conclusions

We explored the post-transcriptional regulatory genetic variants in human tissues and built a genome-wide landscape of genetic variants affecting the percentage of NMD-targeted transcripts (pNMD-QTLs), as well as genetic variants regulating the decay efficiency of NMD-targeted transcripts (dNMD-QTLs). These NMD-QTLs demonstrate a strong association with disease SNPs and exhibit position characteristics. These findings are valuable in understanding post-transcriptional RNA regulation, establishing the etiological roadmap from SNPs to resultant phenotypic traits, and facilitating the research of NMD-based therapeutic strategies for genetic disease.

## Methods

### Data acquisition

For NMD-QTL mapping, we obtained expression data at the transcript level and genotype data from the Genotype-Tissue Expression (GTEx) project (dbGaP accession: phs000424.v7.p2). We grouped transcripts tagged by “nonsense_mediated_decay” in the GENCODE annotation (release 19, GRCh37.p13) as NMD-targeted transcripts for a gene. For disease SNP analysis, we downloaded the curated variant-disease associations from the DisGeNET database (GRCh38) [74]. Genome coordinates from different assembly versions were converted to hg19 through segment_liftover [75]. For miRNA targets, we considered the genome coordinates of targets (conserved or not) for conservative and broad conservative miRNAs obtained in TargetScan [53] which included a total of 257 miRNAs. For RNA binding protein (RBP) binding sites, eCLIP experiments for 120 RBPs from the K562 cell line and 103 RBPs from the HepG2 cell line from ENCODE were used.

### Modeling of NMD-QTLs

Assuming the amount of mRNAs being transcribed is *t*, the percentage of transcribed mRNAs that are subject to NMD is *a*, and the NMD degradation efficiency relative to regular mRNA degradation is (1−*θ*), the amount of NMD-targeted transcripts observed from the RNA-seq experiment is the result of *t* · *a* · *θ*. To apply an additive genetic model, the total level for NMD-targeted transcripts can be written as *T* = *t*_A_ · *α*_A_ · *θ*_A_ + *t*_*a*_ · *α*_*a*_ · *θ*_*a*_, where the subscript A and *a* denote the two alleles for a biallelic SNP. Two scenarios were our major interests: (i) pNMD-QTL where the genetic variant affects the percentage of transcripts subject to NMD (*α*_A_ ≠ *α*_*a*_), (ii) dNMD-QTL where the genetic variant regulates the NMD efficacy (*θ*_A_ ≠ *θ*_*a*_). These two cases can be distinguished from regular eQTLs by examining their effect sizes obtained in our NMD-aware QTL mapping procedure as shown in Fig. 1.

### Identification of NMD-QTLs

The mRNA transcripts of a gene were divided into two categories: NMD-targeted and non-NMD transcripts, based on whether they were annotated by a “nonsense_mediated_decay” tag in the GENCODE annotation. We summarized transcripts level by adding up the transcripts per million (TPM) of all isoforms from the NMD-targeted group and the non-NMD group respectively. We excluded genes with all or none NMD-tagged transcripts. The summarized expressions were normalized within samples by a normalizing factor calculated based on the median of the geometric mean of all genes considered and were then normalized across samples by a rank-based inverse normal transformation. Detailed formulas for normalization can be found in Supplementary Texts.

QTL mappings were performed with FastQTL [76] on both the NMD-targeted and non-NMD groups. According to our modeling, pNMD-QTLs and dNMD-QTLs can be identified by comparing their regulatory effect sizes (Fig. 1). For pNMD-QTLs, the allelic regulatory effect for NMD-targeted transcripts and non-NMD transcripts are in the opposite direction. For dNMD-QTLs, the regulatory effect is only observable for NMD-targeted transcripts but not non-NMD transcripts. In our QTL mapping (NMD-QTLs and splicing-QTLs), similar to the approach in GTEx, variants in the up- or down-stream 1 Mb *cis* window around the TSS were considered. The same covariates used in GTEx eQTL analysis were used. They included genotyping principal components, sequencing platform, sequencing protocol, sex, and a set of covariates identified using the Probabilistic Estimation of Expression Residuals (PEER) method [77]. As in the GTEx eQTL analysis, the permutation mode in FastQTL was used with the setting “--permute 1000 10000” and the false discovery rate (FDR) threshold of 0.05 was applied in our studies.

### Effect size confirmation with quantile regression

RNA-seq quantifications tend to be heavy-tailed and with inflated number of zeros. The effect size estimation in the ordinary linear square fit in FastQTL is sensitive to such non-normality and outliers, even with the applied inverse normal transformation. To reduce false positives and increase the model robustness, for pNMD-QTLs and dNMD-QTLs discovered from FastQTL, we further fit quantile regression models on TPM values without the inverse normal transformation as we did previously [38]. Only pNMD-QTLs and dNMD-QTLs still exhibiting desired effect size properties in quantile regression models were in our final discovery list. Thus, for pNMD-QTLs, opposite signs of effect sizes for NMD-targeted and non-NMD transcripts were observed and the former absolute value was smaller. For dNMD-QTLs, significant effect size was only observed for NMD-targeted transcripts.

### Simulation studies to test our model

To validate our NMD-QTL models, we estimated the precision and recall for identifying pNMD-QTLs and dNMD-QTLs via massive amount of simulations. We simulated the expression levels (total detected transcripts) for NMD-targeted transcripts and non-NMD transcripts by formulas in Fig. 1c for different scenarios. Then we performed QTL calling for both NMD-targeted and non-NMD transcripts through regression models. We declared pNMD-QTL or dNMD-QTL findings according to the desired effect size and directions.

Specifically, we performed three rounds of simulations, each round with different regulatory strength and 10,000 simulated genetic variants. For each genetic variant, n genotypes, X = [x_1_, x_2_, …, x_n_], were simulated with a minor allele frequency of m, where n = 500 and m = 0.05. The total detected transcripts, 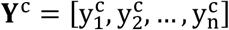, were simulated as a function of genotype, transcript type c (NMD-targeted transcripts or not), and plus a Gaussian random noise: 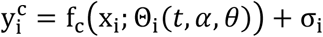.

For the simulated expression and genotype data, we performed a linear regression Y_c_ = *β*_c_X + *ε* for each SNP and inspected the coefficient *β*_c_. If *β*_c_′s were significant (p-value less than a particular threshold) for both NMD-targeted and non-NMD transcripts and were in opposite directions, the SNP was identified as a pNMD-QTL. If *β*_c_ was significant for NMD-targeted transcripts but not for non-NMD transcripts, the SNP was identified as a dNMD-QTL. The detailed parameter settings for simulations with different regulatory strengths can be found in Supplementary Texts.

### Genome-wide NMD-QTL mapping for GTEx tissues

The analysis power for NMD-QTL detection was linearly related to the sample size (Supplementary Fig. 5a, the regression coefficient of the sample size was 28.8 for pNMD-QTLs and 117.0 for dNMD-QTLs), but less sensitive than that for eQTL detection (Supplementary Fig. 5b, regression coefficient: 2938). The testis and whole blood tissues were drastically deviated from this linear trend (Supplementary Fig. 5a), suggesting that the testis had many more NMD-QTLs while the whole blood had many less NMD-QTLs than expected. Such deviations were also observed in the eQTLs mapping (Supplementary Fig. 5b).

### NMD-QTL signature similarity between a tissue pair

To measure the similarity of identified NMD-QTLs between tissues, we calculated s_ij_=n_ij_/(n_i_-n_ij_)/(n_j_/(n-n_j_)) where n_ij_ is the number of NMD-QTLs shared between tissue i and j; n_i_ and n_j_ are the numbers of NMD-QTLs discovered in respective tissues; and n is the total number of nonredundant NMD-QTLs discovered across all tissues. Thus, s_ij_ compares the odds of observing tissue j’s NMD-QTLs in tissue i with the odds of observing tissue j’s NMD-QTLs in all NMD-QTLs. Note that s_ij_ is not necessarily equal to s_ji_, we calculated 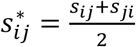 as the final similarity score for NMD-QTL signatures between two tissues. If two tissues share many NMD-QTLs, 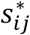 is large.

### Colocalization of NMD-QTLs with disease-associated SNPs

Considering the disease associated variants reported in DisGeNET, we compared how likely the regular eQTLs reported by GTEx and our NMD-QTLs to be colocalized with disease SNPs. We devised a strategy with the combination of a proportion test and a hyper-geometric test to exam the significance of the colocalization between disease SNPs and NMD-QTLs. Here, we pooled NMD-QTLs identified across different tissues. Thus, one NMD-QTL could be counted multiple times if it was identified in multiple tissues. Detail of such two tests can be found in Supplementary Texts.

### Colocalization enrichment score for MeSH

For a disease category represented by a MeSH term (C01-C26, F01-F03), the colocalization enrichment score was calculated as the number of pNMD-QTLs or dNMD-QTLs which were also associated with the disease SNP in the DisGeNET database for the MeSH term, further normalized by the total number of pNMD-QTLs or dNMD-QTLs and the total number disease SNPs annotated by the MeSH term in DisGeNET. The same calculation was performed for eQTL and randomly sampled SNPs.

### Enrichment of NMD-QTLs in the targets of miRNAs and RBPs

miRNA and RBPs were ranked by the number of nonredundant pNMD-QTLs or dNMD-QTLs across all tissues which were located in their target regions and normalized by the total base-pair length of the target regions.

### Splicing-QTL mapping

We identified all cassette exons from the GENCODE annotation. A cassette exon is defined as a non-boundary exon between two other exons and can be either included or skipped to generate two distinct transcripts in alternative splicing. To simplify, we considered transcripts either using or skipping the cassette exon entirely. In other words, transcripts partially overlap with the cassette exons were excluded. To perform splicing-QTL mapping, for a given cassette exon, we used the inclusion ratio of that cassette exon as the quantitative trait. The inclusion ratio of a given cassette exon is calculated as follows:

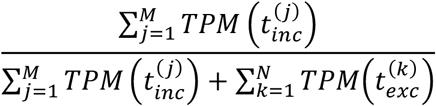

where 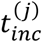 are transcripts including the cassette exon, 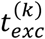 are transcripts excluding the cassette exon, and TPM() is the detected transcripts per million for a transcript. We provide the list of cassette exons and their corresponding transcripts in Supplementary Table 4.

## Declarations

### Ethics approval and consent to participate

Not applicable.

### Consent for publication

Not applicable.

### Availability of data and materials

Original data were obtained from the following online sources:

GTEx Analysis V7: https://gtexportal.org/home/datasets

Gene annotation file: https://www.gencodegenes.org/human/release_19.html

Variant-disease association: https://www.disgenet.org/downloads

Genome assembly conversion tool: https://github.com/baudisgroup/segment-liftover

miRNA targets: http://www.targetscan.org/cgi-bin/targetscan/data_download.vert72.cgi

RBP binding sites (K562 and HepG2):

https://www.encodeproject.org/search/?type=Experiment&control_type!=*&status=released&perturbed=false&assay_title=eCLIP

### Authors’ contributions

BS and LC conceived and designed the work. BS conducted formal analyses. LC supervised the study. BS and LC wrote the manuscript. All authors read and approved the final manuscript.

## Supplementary figures

**Supplementary Figure 1**. NMD-QTL identification results for simulations with various effect sizes. The top panels show the Precision-Recall curves for small effect size (0.05-0.1), medium effect size (0.1-0.15), and large effect size (0.15-0.2). The bottom panels show the p-value distributions for the allelic effect for NMD-targeted transcripts and non-NMD transcripts.

**Supplementary Figure 2**. Ordinal positions of exon and intron intervals in NMD genes. **(a)** The exon or intron interval count per ordinal position. **(b)** The median exon or intron length per ordinal position. **(c)** Distribution of the raw pNMD-QTL or dNMD-QTL count per exon ordinal position. **(d)** Distribution of the raw pNMD-QTL or dNMD-QTL count per intron ordinal position.

**Supplementary Figure 3**. Distances to exon (or intron) boundaries for dNMD-QTLs or eQTLs located on an exon (or intron).

**Supplementary Figure 4**. The distance between the PTC of the NMD-targeted transcripts and the regular stop codon located on the last exon of the non-NMD transcripts. Only exonic positions are counted in the distance calculation.

**Supplementary Figure 5**. Detection power and sample size. **(a)** Positive correlation between the number of NMD-QTLs identified for each tissue and the tissue sample size (i.e., the number of donors). **(b)** The positive correlation observed for GTEx eQTLs as well.

## Supplementary tables

**Supplementary Table 1**. The 43 diseases with significant NMD-QTLs enrichment. p-values for both the proportion test and the hypergeometric test <0.05.

**Supplementary Table 2**. The list of miRNAs and the number of dNMD-QTLs or pNMD-QTLs in their target regions.

**Supplementary Table 3**. The list of RBPs and the number of dNMD-QTLs or pNMD-QTLs in their binding sites in the K562 or the HepG2 cell lines.

**Supplementary Table 4**. The list of cassette exons considered in splicing-QTL mapping.

## References

1. Behm-Ansmant I, Kashima I, Rehwinkel J, Saulière J, Wittkopp N, Izaurralde E. mRNA quality control: an ancient machinery recognizes and degrades mRNAs with nonsense codons. FEBS Lett. 2007;581:2845–53.

2. Bhuvanagiri M, Schlitter AM, Hentze MW, Kulozik AE. NMD: RNA biology meets human genetic medicine. Biochem J. 2010;430:365–77.

3. Huang L, Wilkinson MF. Regulation of nonsense-mediated mRNA decay. Wiley Interdiscip Rev RNA. 2012;3:807–28.

4. Lindeboom RGH, Vermeulen M, Lehner B, Supek F. The impact of nonsense-mediated mRNA decay on genetic disease, gene editing and cancer immunotherapy. Nat Genet. 2019;51:1645–51.

5. Mort M, Ivanov D, Cooper DN, Chuzhanova NA. A meta-analysis of nonsense mutations causing human genetic disease. Hum Mutat. 2008;29:1037–47.

6. Supek F, Lehner B, Lindeboom RGH. To NMD or Not To NMD: Nonsense-Mediated mRNA Decay in Cancer and Other Genetic Diseases. Trends in Genetics. Elsevier; 2021;37:657–68.

7. Mendell JT, Sharifi NA, Meyers JL, Martinez-Murillo F, Dietz HC. Nonsense surveillance regulates expression of diverse classes of mammalian transcripts and mutes genomic noise. Nat Genet. 2004;36:1073–8.

8. Rehwinkel J, Letunic I, Raes J, Bork P, Izaurralde E. Nonsense-mediated mRNA decay factors act in concert to regulate common mRNA targets. RNA. 2005;11:1530–44.

9. Tani H, Imamachi N, Salam KA, Mizutani R, Ijiri K, Irie T, et al. Identification of hundreds of novel UPF1 target transcripts by direct determination of whole transcriptome stability. RNA Biol. 2012;9:1370–9.

10. Weischenfeldt J, Damgaard I, Bryder D, Theilgaard-Mönch K, Thoren LA, Nielsen FC, et al. NMD is essential for hematopoietic stem and progenitor cells and for eliminating by-products of programmed DNA rearrangements. Genes & development. Cold Spring Harbor Lab; 2008;22:1381–96.

11. Zheng S. Alternative splicing and nonsense-mediated mRNA decay enforce neural specific gene expression. International Journal of Developmental Neuroscience. Elsevier; 2016;55:102–8.

12. Lelivelt MJ, Culbertson MR. Yeast Upf proteins required for RNA surveillance affect global expression of the yeast transcriptome. Mol Cell Biol. 1999;19:6710–9.

13. Mitrovich QM, Anderson P. mRNA Surveillance of Expressed Pseudogenes in C. elegans. Current Biology. Elsevier; 2005;15:963–7.

14. Kurosaki T, Maquat LE. Nonsense-mediated mRNA decay in humans at a glance. J Cell Sci. 2016;129:461–7.

15. Lykke-Andersen S, Jensen TH. Nonsense-mediated mRNA decay: an intricate machinery that shapes transcriptomes. Nat Rev Mol Cell Biol. 2015;16:665–77.

16. Ottens F, Gehring NH. Physiological and pathophysiological role of nonsense-mediated mRNA decay. Pflugers Arch. 2016;468:1013–28.

17. Raimondeau E, Bufton JC, Schaffitzel C. New insights into the interplay between the translation machinery and nonsense-mediated mRNA decay factors. Biochem Soc Trans. 2018;46:503–12.

18. Han X, Wei Y, Wang H, Wang F, Ju Z, Li T. Nonsense-mediated mRNA decay: a “nonsense” pathway makes sense in stem cell biology. Nucleic Acids Res. 2018;46:1038–51.

19. Shum EY, Jones SH, Shao A, Dumdie J, Krause MD, Chan W-K, et al. The Antagonistic Gene Paralogs Upf3a and Upf3b Govern Nonsense-Mediated RNA Decay. Cell. 2016;165:382–95.

20. Bao J, Tang C, Yuan S, Porse BT, Yan W. UPF2, a nonsense-mediated mRNA decay factor, is required for prepubertal Sertoli cell development and male fertility by ensuring fidelity of the transcriptome. Development. 2015;142:352–62.

21. Bao J, Vitting-Seerup K, Waage J, Tang C, Ge Y, Porse BT, et al. UPF2-Dependent Nonsense-Mediated mRNA Decay Pathway Is Essential for Spermatogenesis by Selectively Eliminating Longer 3’UTR Transcripts. PLoS Genet. 2016;12:e1005863.

22. Kurosaki T, Sakano H, Pröschel C, Wheeler J, Hewko A, Maquat LE. NMD abnormalities during brain development in the Fmr1-knockout mouse model of fragile X syndrome. Genome Biology. 2021;22:317.

23. Li Z, Vuong JK, Zhang M, Stork C, Zheng S. Inhibition of nonsense-mediated RNA decay by ER stress. RNA. 2017;23:378–94.

24. Zhao J, Li Z, Puri R, Liu K, Nunez I, Chen L, et al. Molecular profiling of individual FDA-approved clinical drugs identifies modulators of nonsense-mediated mRNA decay. Mol Ther Nucleic Acids. 2022;27:304–18.

25. Gudikote JP, Imam JS, Garcia RF, Wilkinson MF. RNA splicing promotes translation and RNA surveillance. Nat Struct Mol Biol. 2005;12:801–9.

26. Usuki F, Yamashita A, Higuchi I, Ohnishi T, Shiraishi T, Osame M, et al. Inhibition of nonsense-mediated mRNA decay rescues the phenotype in Ullrich’s disease. Annals of Neurology. 2004;55:740–4.

27. Zetoune AB, Fontanière S, Magnin D, Anczuków O, Buisson M, Zhang CX, et al. Comparison of nonsense-mediated mRNA decay efficiency in various murine tissues. BMC Genetics. 2008;9:83.

28. Huang L, Lou C-H, Chan W, Shum EY, Shao A, Stone E, et al. RNA Homeostasis Governed by Cell Type-Specific and Branched Feedback Loops Acting on NMD. Molecular Cell. 2011;43:950–61.

29. Miller JN, Pearce DA. Nonsense-mediated decay in genetic disease: friend or foe? Mutat Res Rev Mutat Res. 2014;762:52–64.

30. Consortium TGte. The Genotype-Tissue Expression (GTEx) pilot analysis: Multitissue gene regulation in humans. Science. American Association for the Advancement of Science; 2015;348:648–60.

31. GTEx Consortium, Laboratory, Data Analysis &Coordinating Center (LDACC)—Analysis Working Group, Statistical Methods groups—Analysis Working Group, Enhancing GTEx (eGTEx) groups, NIH Common Fund, NIH/NCI, et al. Genetic effects on gene expression across human tissues. Nature. 2017;550:204–13.

32. Wu L, Candille SI, Choi Y, Xie D, Li-Pook-Than J, Tang H, et al. Variation and Genetic Control of Protein Abundance in Humans. Nature. 2013;499:79–82.

33. Hsiao Y-HE, Bahn JH, Lin X, Chan T-M, Wang R, Xiao X. Alternative splicing modulated by genetic variants demonstrates accelerated evolution regulated by highly conserved proteins. Genome Res. 2016;26:440–50.

34. Zhang H, Dou S, He F, Luo J, Wei L, Lu J. Genome-wide maps of ribosomal occupancy provide insights into adaptive evolution and regulatory roles of uORFs during Drosophila development. PLOS Biology. Public Library of Science; 2018;16:e2003903.

35. MacArthur DG, Tyler-Smith C. Loss-of-function variants in the genomes of healthy humans. Hum Mol Genet. 2010;19:R125–30.

36. Rivas MA, Pirinen M, Conrad DF, Lek M, Tsang EK, Karczewski KJ, et al. Impact of predicted protein-truncating genetic variants on the human transcriptome. Science. 2015;348:666–9.

37. Teran NA, Nachun DC, Eulalio T, Ferraro NM, Smail C, Rivas MA, et al. Nonsense-mediated decay is highly stable across individuals and tissues. Am J Hum Genet. 2021;108:1401–8.

38. Sun B, Chen L. Quantile regression for challenging cases of eQTL mapping. Briefings in Bioinformatics. 2020;21:1756–65.

39. Nguyen LS, Wilkinson MF, Gecz J. Nonsense-mediated mRNA decay: inter-individual variability and human disease. Neurosci Biobehav Rev. 2014;46 Pt 2:175–86.

40. Khajavi M, Inoue K, Lupski JR. Nonsense-mediated mRNA decay modulates clinical outcome of genetic disease. Eur J Hum Genet. 2006;14:1074–81.

41. Piñero J, Bravo À, Queralt-Rosinach N, Gutiérrez-Sacristán A, Deu-Pons J, Centeno E, et al. DisGeNET: a comprehensive platform integrating information on human disease-associated genes and variants. Nucleic Acids Res. 2017;45:D833–9.

42. Nguyen LS, Kim H-G, Rosenfeld JA, Shen Y, Gusella JF, Lacassie Y, et al. Contribution of copy number variants involving nonsense-mediated mRNA decay pathway genes to neurodevelopmental disorders. Hum Mol Genet. 2013;22:1816–25.

43. Tarpey PS, Raymond FL, Nguyen LS, Rodriguez J, Hackett A, Vandeleur L, et al. Mutations in UPF3B, a member of the nonsense-mediated mRNA decay complex, cause syndromic and nonsyndromic mental retardation. Nat Genet. 2007;39:1127–33.

44. Addington AM, Gauthier J, Piton A, Hamdan FF, Raymond A, Gogtay N, et al. A novel frameshift mutation in UPF3B identified in brothers affected with childhood onset schizophrenia and autism spectrum disorders. Mol Psychiatry. 2011;16:238–9.

45. Laumonnier F, Shoubridge C, Antar C, Nguyen LS, Van Esch H, Kleefstra T, et al. Mutations of the UPF3B gene, which encodes a protein widely expressed in neurons, are associated with nonspecific mental retardation with or without autism. Mol Psychiatry. 2010;15:767–76.

46. Colombo M, Karousis ED, Bourquin J, Bruggmann R, Mühlemann O. Transcriptome-wide identification of NMD-targeted human mRNAs reveals extensive redundancy between SMG6- and SMG7-mediated degradation pathways. RNA. 2017;23:189–201.

47. Le Hir H, Gatfield D, Izaurralde E, Moore MJ. The exon-exon junction complex provides a binding platform for factors involved in mRNA export and nonsense-mediated mRNA decay. EMBO J. 2001;20:4987–97.

48. Brogna S, Wen J. Nonsense-mediated mRNA decay (NMD) mechanisms. Nat Struct Mol Biol. 2009;16:107–13.

49. Bartel DP. Metazoan MicroRNAs. Cell. 2018;173:20–51.

50. Bartel DP. MicroRNAs: target recognition and regulatory functions. Cell. 2009;136:215–33.

51. Hogan DJ, Riordan DP, Gerber AP, Herschlag D, Brown PO. Diverse RNA-binding proteins interact with functionally related sets of RNAs, suggesting an extensive regulatory system. PLoS Biol. 2008;6:e255.

52. Lunde BM, Moore C, Varani G. RNA-binding proteins: modular design for efficient function. Nat Rev Mol Cell Biol. 2007;8:479–90.

53. Agarwal V, Bell GW, Nam J-W, Bartel DP. Predicting effective microRNA target sites in mammalian mRNAs. Izaurralde E, editor. eLife. eLife Sciences Publications, Ltd; 2015;4:e05005.

54. An Integrated Encyclopedia of DNA Elements in the Human Genome. Nature. 2012;489:57–74.

55. Zheng S, Black DL. Alternative pre-mRNA splicing in neurons: growing up and extending its reach. Trends Genet. 2013;29:442–8.

56. Zheng S. Alternative splicing and nonsense-mediated mRNA decay enforce neural specific gene expression. Int J Dev Neurosci. 2016;55:102–8.

57. Giorgi C, Yeo GW, Stone ME, Katz DB, Burge C, Turrigiano G, et al. The EJC factor eIF4AIII modulates synaptic strength and neuronal protein expression. Cell. 2007;130:179–91.

58. Yap K, Makeyev EV. Regulation of gene expression in mammalian nervous system through alternative pre-mRNA splicing coupled with RNA quality control mechanisms. Mol Cell Neurosci. 2013;56:420–8.

59. Zheng S, Gray EE, Chawla G, Porse BT, O’Dell TJ, Black DL. PSD-95 is posttranscriptionally repressed during early neural development by PTBP1 and PTBP2. Nat Neurosci. 2012;15:381–8, S1.

60. Aguet F, Brown AA, Castel SE, Davis JR, He Y, Jo B, et al. Genetic effects on gene expression across human tissues. Nature. 2017;550:204–13.

61. Morley M, Molony CM, Weber TM, Devlin JL, Ewens KG, Spielman RS, et al. Genetic analysis of genome-wide variation in human gene expression. Nature. 2004;430:743–7.

62. Stranger BE, Forrest MS, Dunning M, Ingle CE, Beazley C, Thorne N, et al. Relative impact of nucleotide and copy number variation on gene expression phenotypes. Science. 2007;315:848–53.

63. Montgomery SB, Sammeth M, Gutierrez-Arcelus M, Lach RP, Ingle C, Nisbett J, et al. Transcriptome genetics using second generation sequencing in a Caucasian population. Nature. 2010;464:773–7.

64. Pickrell JK, Marioni JC, Pai AA, Degner JF, Engelhardt BE, Nkadori E, et al. Understanding mechanisms underlying human gene expression variation with RNA sequencing. Nature. 2010;464:768–72.

65. Garrido-Martín D, Borsari B, Calvo M, Reverter F, Guigó R. Identification and analysis of splicing quantitative trait loci across multiple tissues in the human genome. Nat Commun. 2021;12:727.

66. Melé M, Ferreira PG, Reverter F, DeLuca DS, Monlong J, Sammeth M, et al. Human genomics. The human transcriptome across tissues and individuals. Science. 2015;348:660–5.

67. Lonsdale J, Thomas J, Salvatore M, Phillips R, Lo E, Shad S, et al. The Genotype-Tissue Expression (GTEx) project. Nature Genetics. 2013;45:580–5.

68. Keeling KM, Bedwell DM. Suppression of nonsense mutations as a therapeutic approach to treat genetic diseases. Wiley Interdiscip Rev RNA. 2011;2:837–52.

69. Sartor F, Anderson J, McCaig C, Miedzybrodzka Z, Müller B. Mutation of genes controlling mRNA metabolism and protein synthesis predisposes to neurodevelopmental disorders. Biochem Soc Trans. 2015;43:1259–65.

70. Balistreri G, Bognanni C, Mühlemann O. Virus Escape and Manipulation of Cellular Nonsense-Mediated mRNA Decay. Viruses. 2017;9:E24.

71. Linde L, Boelz S, Nissim-Rafinia M, Oren YS, Wilschanski M, Yaacov Y, et al. Nonsense-mediated mRNA decay affects nonsense transcript levels and governs response of cystic fibrosis patients to gentamicin. J Clin Invest. 2007;117:683–92.

72. Keeling KM. Nonsense Suppression as an Approach to Treat Lysosomal Storage Diseases. Diseases. 2016;4:32.

73. Nomakuchi TT, Rigo F, Aznarez I, Krainer AR. Antisense oligonucleotide-directed inhibition of nonsense-mediated mRNA decay. Nat Biotechnol. 2016;34:164–6.

74. Piñero J, Ramírez-Anguita JM, Saüch-Pitarch J, Ronzano F, Centeno E, Sanz F, et al. The DisGeNET knowledge platform for disease genomics: 2019 update. Nucleic Acids Research. 2020;48:D845–55.

75. Gao B, Huang Q, Baudis M. segment_liftover : a Python tool to convert segments between genome assemblies. F1000Res. 2018;7:319.

76. Ongen H, Buil A, Brown AA, Dermitzakis ET, Delaneau O. Fast and efficient QTL mapper for thousands of molecular phenotypes. Bioinformatics. 2016;32:1479–85.

77. Stegle O, Parts L, Durbin R, Winn J. A Bayesian Framework to Account for Complex Non-Genetic Factors in Gene Expression Levels Greatly Increases Power in eQTL Studies. Regev A, editor. PLoS Computational Biology. 2010;6:e1000770.

