## Supplementary material for "Mapping genetic variants for nonsense-mediated mRNA decay regulation across human tissues": Supp_F2.pdf

(a) exon/intron count by ordinal position

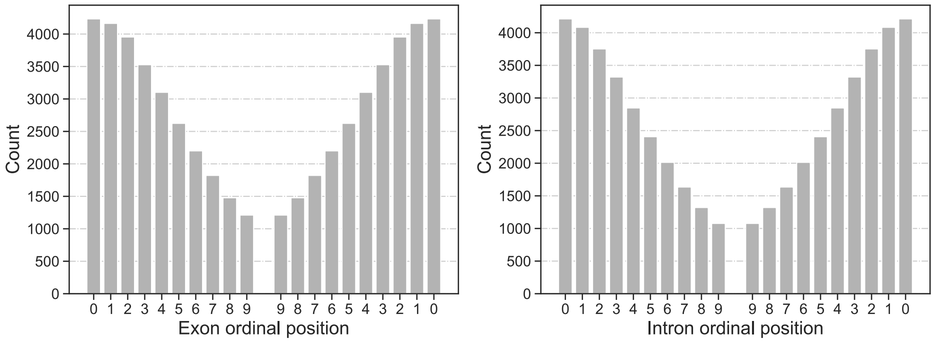


(b) Median exon/intron length by ordinal position

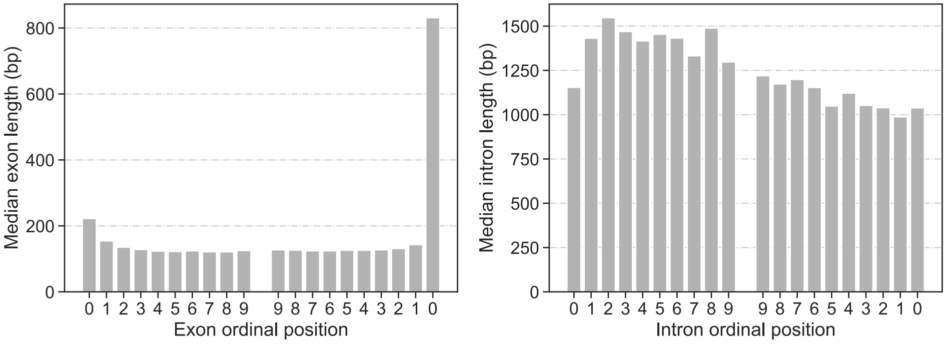


(c) Raw NMD-QTL count by exon ordinal position

(d) Raw NMD-QTL count by intron ordinal position

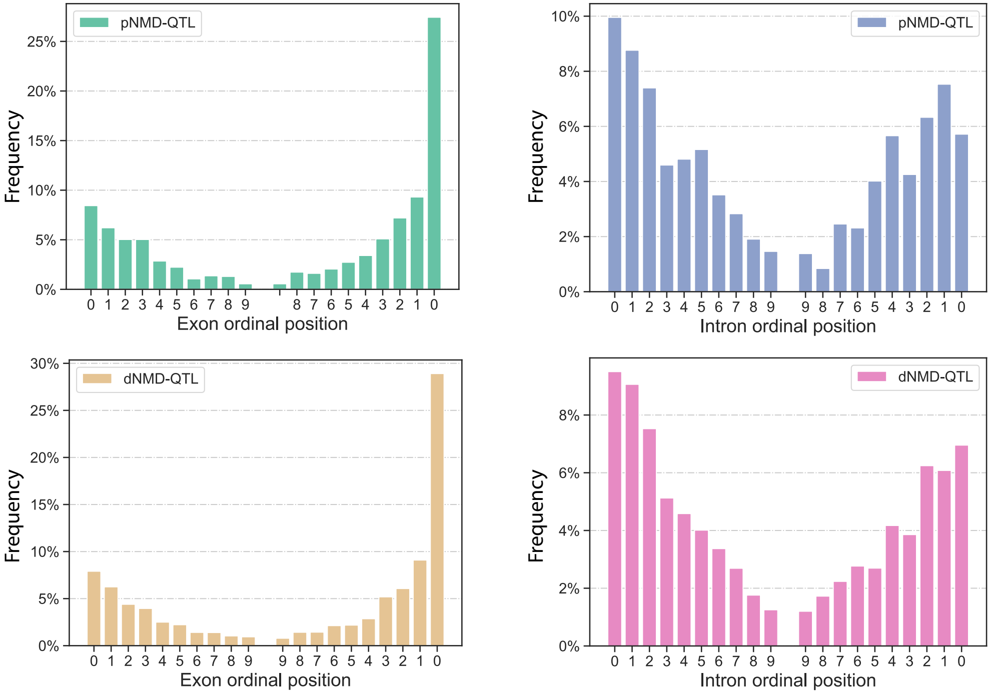
