## Supplementary material for "Mapping genetic variants for nonsense-mediated mRNA decay regulation across human tissues": SupplementaryTexts.pdf

### Supplementary Texts

**Expression normalization.** We performed NMD-QTL mapping for 48 tissues from GTEx with the sample size ≥ 70. For a given tissue, the NMD expression and non-NMD expression were obtained by the following procedures:

- Lowly expressed genes were filtered out. Only genes with TPM > 0.1 in at least 10 samples were selected for further analysis.
- The filtered genes expression was summarized into NMD expression and non-NMD expression.
- Since NMD expression could be sparse, only genes with the percent of zeros less than 40% in NMD expression were kept.
- Phenotypic traits (NMD expression and non-NMD expression) were normalized based on a normalization factor. Suppose the total number of genes is G and the total number of individuals is K, the normalization factor for each sample k was calculated based on all expressed genes as follows:
  - First a geometric mean was calculated for each gene g across samples:

$$\mu^{(g)} = \left( \prod_{k:TPM_g^{(k)} \neq 0} TPM_g^{(k)} \right)^{1/\sum_k I\left(TPM_g^{(k)} \neq 0\right)}$$

  - Then, the normalization factor for each individual k is obtained as the median fold-change of expressed genes compared to this geometric mean: $\delta^{(k)} = \mathrm{median}\left(\frac{\mathrm{TPM}_g^{(k)}}{\mu^{(g)}},\ \mathrm{for}\ g \in G\right)$
  - then the NMD expression and non-NMD expression were normalized by this normalization factor, more formally, normalized TPM for every gene g and sample k is then given by: $TPM'^{(k)}_g = \frac{TPM_g^{(k)}}{\delta^{(k)}}$.

- Finally, NMD expression and non-NMD expression were further normalized across samples by the rank-based inverse normal transform used in FastQTL for NMD-QTL mapping.

Python code snippet for the normalization logic and the inverse normal transform is shown below.

```
import pandas as pd
import scipy.stats as ss
from scipy.stats.mstats import gmean
def rank_INT(x, c=3./8):
   """Rank-based Inverse Normal Transform.
     Ties share the same value after transformation."""
   n = len(x)
   r = ss.rankdata(x, method='average')
   return ss.norm.ppf((r - c) / (n - 2 * c + 1))

# normalization factors
gene_tpm = pd.read_csv(os.path.join(base_path, 'GTEx_Analysis_2016-01-
15_v7_RNASeQCv1.1.8_gene_tpm.gct.gz'),
            skiprows=2, sep='\t', usecols=tissue_cols[2:])
gene_tpm = gene_tpm[(gene_tpm > .1).sum(axis=1) >= 10]  # filter out lowly expressed genes
genes_gmean = gene_tpm.apply(lambda row: gmean(row[row != 0]), axis=1)
norm_factors = gene_tpm.divide(genes_gmean, axis=0).median(axis=0)
```

**Detailed parameter settings used in simulations.** Without any prior knowledge on how the NMD effect should be, parameters $\Theta(t\ , \alpha\ , \theta\ )$ were generated by first drawing $\alpha_A$ and $\theta_A$ from a uniform distribution U(0.2, 0.8), and then obtaining $\alpha_a$, and $\theta_a$ by ensuring the effect size $|\alpha_A - \alpha_a|$ and $|\theta_A - \theta_a|$ belonging to the desired range. For cases assuming $t_A$=$t_a$=t we chose t = 4, and for cases assuming $t_A \neq t_a$, we drew both $t_A \sim U(3,5)$ and $t_a \sim U(3,5)$, and then applied a Gaussian noise term. We drew the Gaussian noise term $\sigma \sim N(0,1.5)$ and truncated $y_i$ to be non-negative to mimic the real data.

**Hyper-geometric test used in disease SNP colocalization study for NMD-QTLs.** For a given disease $d_i$, let $N$ be the total number of NMD-QTLs discovered, $M$ be the universal set of SNPs (here we assume it is the number of all SNPs tested in GTEx), let $n$ be the number of markers reported in DisGeNET for $d_i$, and $x$ be the number of NMD-QTLs that were indeed the disease marker for $d_i$. Then the p-value for the test is given by

$$p = \mathbf{Pr}(X \geq x - 1)$$

where X follows a hypergeometric distribution of which the probability mass function is defined as

$$p(x, M, n, N) = \frac{\binom{n}{x}\binom{M-n}{N-x}}{\binom{M}{N}}.$$

**Proportion test used in disease SNP colocalization study for NMD-QTLs.** For a given disease $d_i$, four statistics were gathered as follows. The alternative hypothesis was $H_A: \frac{x_1}{y_1} > \frac{x_2}{y_2}$.

- $x_1$: number of NMD-QTLs reported to be associated with the $d_i$
- $y_1$: total number of NMD-QTLs reported to be associated diseases
- $x_2$: number of eQTLs reported to be associated with the $d_i$
- $y_2$: total number of eQTLs reported to be associated diseases
