## Supplementary figures and images for "Mapping genetic variants for nonsense-mediated mRNA decay regulation across human tissues"

### Supp_F1.pdf

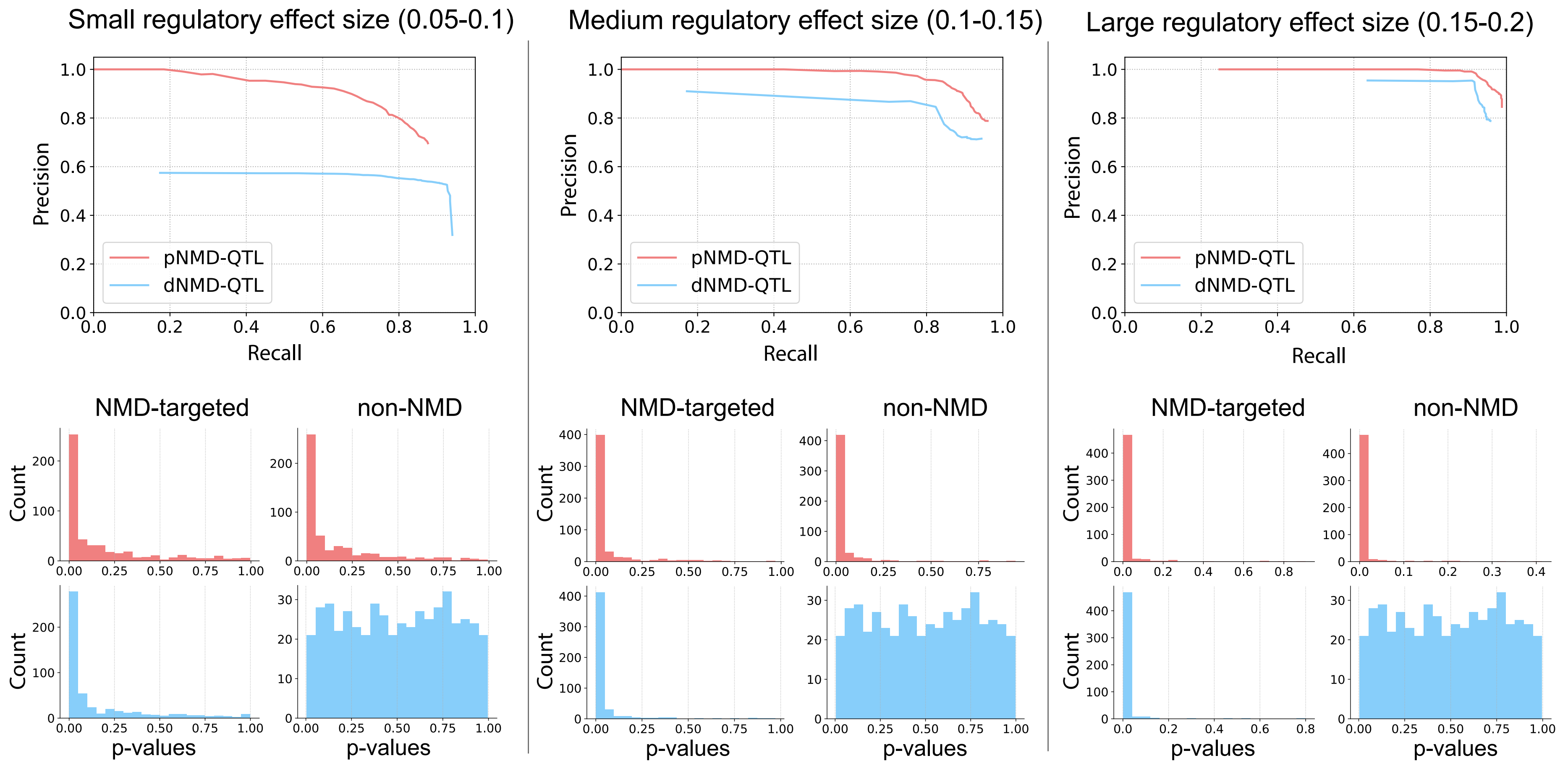

### Supp_F3.pdf

#### (a) Distance to exon boundary (bp)

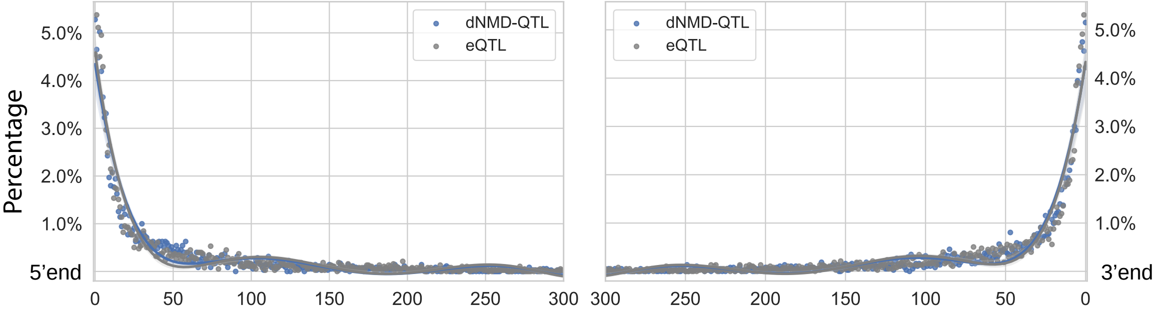


#### (b) Distance to inton boundary (bp)

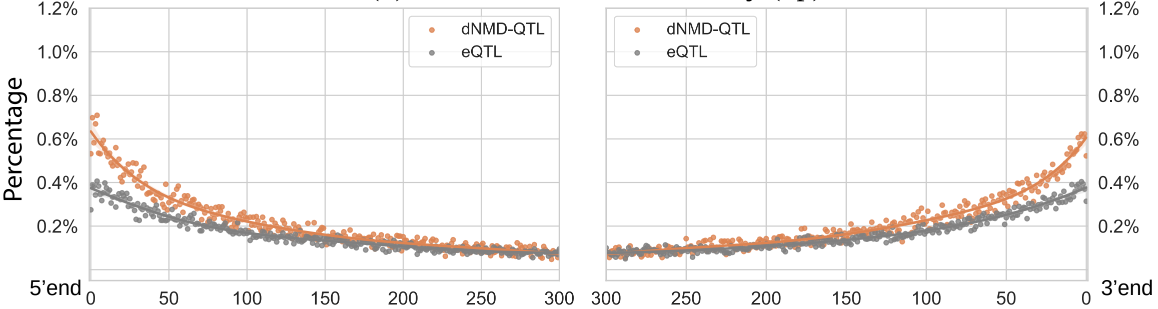

### Supp_F4.pdf

##### distance of PTC to the regular stop codon

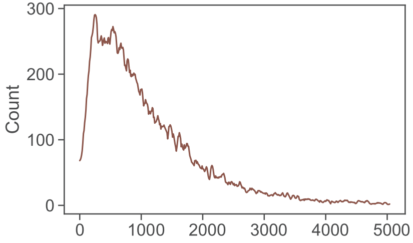

### Supp_F5.pdf

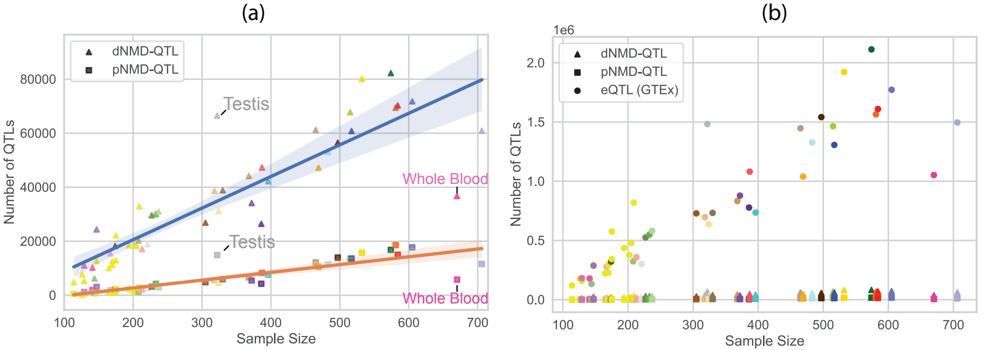
